# Developmental factors drive the compartmentalised and discontinuous maturation of the small intestinal epithelium during the early postnatal period

**DOI:** 10.1101/2025.05.15.654215

**Authors:** Johannes Schöneich, Aline Dupont, Stefan Schlößer, Matthias A. Schmitz, Isabel Richter, Narasimha Murthy Keshava Prasad Gubbi, Mingbo Cheng, Tiago Maié, Sofia Jäverfelt, Fabian Imdahl, Christophe Toussaint, Antoine-Emmanuel Saliba, Christoph Kuppe, Thaher Pelaseyed, Ivan G. Costa, Mathias W. Hornef

**Affiliations:** Institute of Medical Microbiology, RWTH Aachen University Hospital, 52074 Aachen, Germany; Institute for Computational Genomics, RWTH Aachen University Hospital, 52074 Aachen, Germany; Department of Medical Biochemistry and Cell Biology, Institute of Biomedicine, University of Gothenburg, 405 30 Gothenburg, Sweden; Helmholtz Institute for RNA-based Infection Research (HIRI), Helmholtz-Center for Infection Research (HZI), Würzburg, Germany; Department of Nephrology, Faculty of Medicine, RWTH Aachen University, Aachen, Germany

**Author notes:** Contributed equally (Johannes Schöneich, Aline Dupont). Corresponding author: Mathias W. Hornef, M.D., Institute of Medical Microbiology; RWTH Aachen University Hospital; Pauwelsstr. 30, D-52074 Aachen, Germany. Phone: *49 241 80 89510; Fax: *49 241 80 82483;.

**Keywords:** small intestinal epithelium, neonate, postnatal maturation, crypt-villus axis, Salmonella

## Abstract

The temporal, spatial and cellular diversity of the small intestinal epithelium during the postnatal period, a critical time window that accompanies the transition from placental energy supply to enteral feeding, the establishment of the enteric microbiota and postnatal immune maturation, has not been systematically investigated. Here, we used laser capture microdissection and bulk RNA-Seq, proteomics, and single cell RNA-Seq to analyse the total, organ site, crypt- and villus-specific intestinal epithelium of specific pathogen-free, germ-free and *Salmonella*-infected mice during the postnatal period. We identified key temporal and organ-site specific expression patterns that revealed a weak effect of the microbiota but strong influence of developmental regulators during early life. We also determined age-dependent signalling pathway and transcription factor activity and characterised age- and cell type-specific developmental trajectories revealing a distinct compartmentalised maturation process along the proximal-to-distal length and crypt-villus axis and a discontinuous appearance of goblet/Paneth cell and absorptive enterocyte transcriptional profiles. Finally, we described the cell type-specific response to neonatal enteric infection. Taken together, our findings identify the epithelium as an integral element in the maturation of postnatal mucosal tissues and in the establishment of host-microbe homeostasis.

## Introduction

The small intestinal epithelium facilitates the digestion and absorption of dietary nutrients, forms a physicochemical mucosal barrier that restricts the luminal enteric microbiota, and represents the first line of defence in the event of infection. The wide range of functions of the small intestine is reflected by the broad diversity of epithelial cell types such as stem and transit amplifying (TA) cells, absorptive enterocytes, secretory Paneth, goblet, tuft and enteroendocrine cells (EEC), and M cells, all in turn composed of different subtypes and differentiation stages ^1–5^. This cellular diversity is regulated by a complex network of transcriptional and non-transcriptional endogenous factors ^6–13^. In addition, intestinal epithelial cells actively interact with the microbiota and the underlying immune and stroma cells ^14–19^. While intestinal epithelial cell heterogeneity has been characterised in adult mice, as well as in the paediatric and adult human host ^2,3,20,21^, much less is known about the early postnatal period. This period, however, is particularly critical since it facilitates the transition from placental energy supply to enteral feeding and the concomitant establishment of the enteric microbiota and maturation of the mucosal immune system, critical to the maintenance of the host-microbe homeostasis. In addition, the murine neonatal intestinal epithelium differs significantly from that of the adult animal ^1,20^. Crypts and Paneth cells both appear postnatally only around the second week of life ^22,23^. In the absence of crypts, it also lacks defined TA cells, continuous epithelial cell proliferation, and crypt-villus migration ^24^. On the other hand, foetal-type enterocytes that engulf luminal material for intraepithelial processing and the efferocytosis of damaged epithelial cells by neighbouring enterocytes are restricted to the neonatal period ^25,26^. Significant differences in the transcription of metabolic enzymes, differentiation factors, innate immune sensing and antimicrobial host defence molecules have been observed between neonatal and adult mice ^6,7,27–30^. These differences, controlled in part by master regulators of developmental epithelial gene expression such as BLIMP1 (encoded by *Prdm1*) or MAFB, may as well result from microbial signals ^2,7,8,14^.

The aim of the present study was to characterise the multidimensional changes of the small intestinal epithelium during the neonate-to-adult transition. This period of life has attracted particular attention due to the strongly enhanced infection-associated mortality. Furthermore, epidemiological and experimental evidence have suggested that exposure to exogenous factors during this early period can have lasting consequences and determine long-term health ^31,32^. State-of-the-art methods were used to characterise the global, organ-, site- and cell type-specific transcriptional and/or proteomic profile of the gut epithelium of healthy specific pathogen-free (SPF) and germ-free (GF) mice at different time points after birth as well as mice with neonatal enteric infection (Fig. 1a). Our results represent an important new resource for the community and expand our understanding of the age-dependent changes occurring during this transitional period. Characterisation of the influence of environmental factors may help to identify new strategies to promote health at early age and reduce childhood mortality.

**Figure 1:**
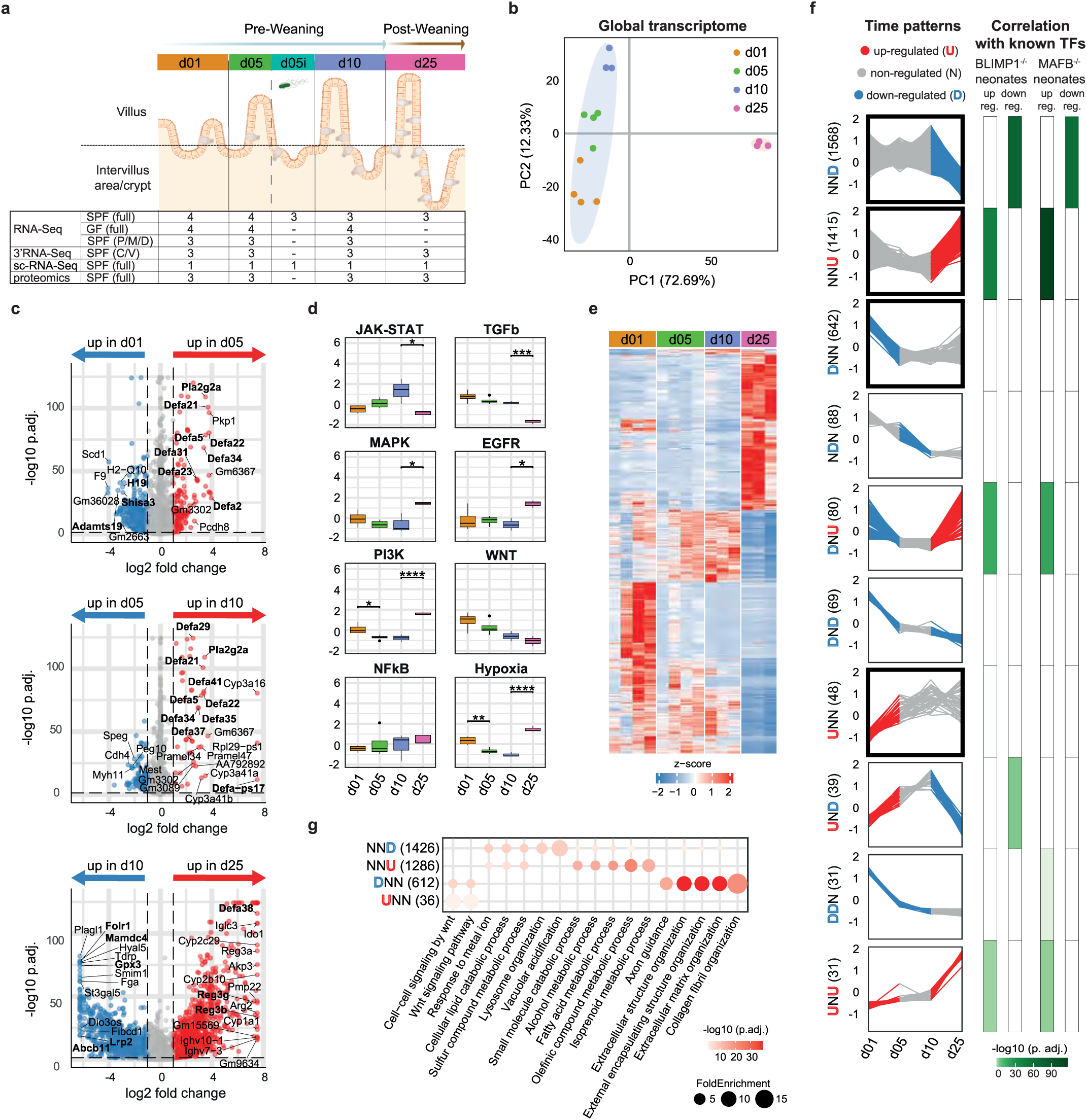
The global intestinal epithelial transcriptome during postnatal development. **(a)** Schematic representation of the murine small intestine development during the early postnatal period (top) and table summarising the different samples (number of biological replicates indicated) analysed in this study (bottom). P: proximal, M: medial, D: distal, C: crypt, V: villus. Figure created using Biorender.com. **(b)** PCA plot representing the full small intestinal epithelial transcriptome of 1-day-old (orange, n=4), 5-day-old (green, n=4), 10-day-old (blue, n=3) and 25-day-old (pink, n=3) mice. Ellipses were estimated using the Khachiyan algorithm and show the distribution of the pre-weaning (light blue) and post- weaning (light pink) samples. **(c)** Volcano plots showing differential gene expression between 1- and 5-day-old (top), 5- and 10-day-old (middle) and 10- and 25-day-old (bottom) samples. Blue and red dots indicate genes statistically (p_adj_<0.05) upregulated (|log2FC| > 1) in the younger and older animals, respectively. Grey dots indicate non-significantly and/or non-differentially regulated genes. **(d)** Box plots showing PROGENy pathway analysis at the different ages. Colour code as in Fig. 1b. **(e)** Heatmap depicting the expression level of all genes differentially regulated between either 1- and 5-day-old, 5- and 10-day-old and/or 10 and 25-day-old animals. **(f)** Ten most abundant temporal postnatal gene expression patterns as identified by temporal DE analysis (U, upregulated, red; N, non-regulated, grey; D, downregulated, blue) and their association with reported BLIMP/MAFB gene regulation. The number of DE genes for each pattern is indicated in brackets. **(g)** Dot plot showing the top 5 enriched GO terms (by adjusted p-value) for the differentially expressed genes identified as belonging to the NND, NNU, DNN and UNN expression patterns. The number of DE genes associated with a GO term for each pattern is indicated in brackets. Dataset available under GSE283495.

## Material & Methods

### Animals

All animal work was performed according to the regulation specified by the Federation for Laboratory Animal Science Associations (FELASA) and the German Society of Laboratory Animal Science (GV- SOLAS, http://www.gv-solas.de). C57BL/6J SPF mice were bred and maintained under specific pathogen-free conditions in ventilated cages, with 12-hour light/dark cycles, and ad libitum access to food and water at the institute of laboratory animal science at the RWTH University Hospital Aachen. GF mice were generously provided by Tom Clavel and Susan A. Jennings (RWTH University Hospital, Aachen). Experimental animal work was approved by the North Rhine-Westphalia Office of Nature, Environment and Consumer Protection (LANUV, approvals 81-02.04.2017.A397). 1-day-old C57BL/6J SPF newborn mice were orally infected with 1µl PBS containing 100-300 log-phase *S.* Typhimurium (ATCC14028). At 4 days post-infection, all animals were sacrificed by decapitation and their small intestines removed and processed as described below.

### Sample preparation for bulk RNA-Seq and proteomics

Intestinal epithelial cells were isolated as previously described ^27^. Briefly, full small intestines or parts of the small intestines (the proximal part corresponding to the proximal 5 to 20% of the full small intestine, the medial part to the 37.5 to 62.5% and the distal part to the 80 to 95%) were cut into small pieces, incubated at 37°C in 30 mM EDTA for 10 minutes and shaken for 20 seconds. Detached epithelial cells were then filtered through a 100µm cell strainer, washed with PBS and pelleted by centrifugation. Pellets were then either resuspended in TRIzol reagent (RNA-Seq) or snap-frozen in liquid nitrogen (proteomics) and stored at -80°C.

### Bulk RNA-Seq

RNA was subsequently isolated following manufacturer’s instructions and cDNA libraries were prepared using NEBNext ultra II directional RNA library prep kit for Illumina with the NEBNext poly(A) mRNA magnetic isolation module (New England Biolabs) and sequenced in paired end mode on a NextSeq 500 or a NovaSeq 6000 sequencer (Illumina) at the Genomics Facility, a core facility of the Interdisciplinary Centre for Clinical Research (IZKF) Aachen within the Faculty of Medicine at RWTH Aachen University. Each sample corresponded to one given animals, except for day 1 samples for which the cells isolated from the small intestine of two 1-day-old animals were pooled.

### Proteomic analysis

Each sample corresponded to one given animal, except for day 1 samples for which the cells isolated from the small intestine of three 1-day-old animals were pooled. For the sample preparation, isolated epithelial cells were lysed by addition of 150 µL lysis buffer (4% SDS, 100mM Tris-HCl pH 7.5, 100mM DTT) and heated for 5 min at 95°C. Cell lysates were sonicated for 10 s, centrifuged at 16,000 x g in a tabletop centrifuge at RT and supernatant added onto 10 kDa cutoff filters (#OD010C33, PALL). Protein concentrations were measured at 280 nm using Nanodrop (Thermo Fisher Scientific). 250 µg of protein of each sample was alkylated and digested using filter-aided sample preparation ^33^. Every sample was incubated with 0.45 µg trypsin at 37°C overnight apart from the pooled sample of three 1-day old animals where 0.65 µg trypsin was used. Peptide concentration after elution was measured at 280 nm using NanoDrop and peptides cleaned with StageTip C18 columns prior to mass-spectrometry (MS) analysis ^34^.

Nano liquid chromatography (LC)-MS/MS was performed on a Q-Exactive HF mass-spectrometer (Thermo Fischer Scientific), connected with an EASY-nLC 1000 system (Thermo Fischer Scientific) through a nanoelectrospray ion source. Peptides were loaded on a reverse-phase column (150 mm^3^ 0.075 mm inner diameter, New Objective, New Objective, Woburn, MA) packed in-house with Reprosil-Pur C18-AQ 3 mm particles (Dr. Maisch, Ammerbuch, Germany). Peptides were separated with a 230-minute gradient: from 3 to 25% B in 175 min, 25 to 45% B in 30 min, 45 to 100% B in 5 min, followed 20 min wash with 100% of B (A: 0.1% formic acid, B: 0.1% formic acid/80% acetonitrile) using a flow rate of 250 nl/min. Q-Exactive HF was operated at 250°C capillary temperature and 2.0 kV spray voltage. Full mass spectra were acquired in the Orbitrap mass analyzer over a mass range from m/z 350 to 1600 with resolution of 60 000 (m/z 200) after accumulation of ions to a 3e6 target value based on predictive AGC from the previous full scan. Twelve most intense peaks with a charge state ≥2 were fragmented in the HCD collision cell with normalized collision energy of 27%, and tandem mass spectrum was acquired in the Orbitrap mass analyzer with resolution of 15 000 after accumulation of ions to a 1e5 target value. Dynamic exclusion was set to 30 s. The maximum allowed ion accumulation times were 20 ms for full MS scans and 50 ms for tandem mass spectrum.

### Sample preparation for single cell RNA-Seq

Here, single intestinal epithelial cells were isolated using a protocol adapted from Haber et al. ^2^. Longitudinally cut-opened small intestines were placed in a 50 mL tube filled with PBS, shortly vortexed to remove the luminal content and further cut into 2- to 5-mm long pieces. The intestinal pieces were then transferred into a 50 mL tube prefilled with 30 mL room-tempered RPMI supplemented with 5% foetal calf serum (FCS) and 2 mM EDTA. Next, the tube was placed on a shaker set at 100 rpm at room temperature. After 15 minutes incubation, single epithelial cells were filtered through a 100 µm cell strainer, pelleted by centrifugation at 300xg at 4°C. The pellet was resuspended in 1mL PBS containing 5% FCS and 0,01 mM EDTA and kept on ice. The remaining tissue pieces were transferred into a new 50 mL tubes containing 30 mL room-tempered RPMI supplemented with 5% foetal calf serum (FCS) and 2mM EDTA and the cell isolation process was repeated three more times. The first fraction collected was discarded, while the epithelial cells contained in the three last fractions were pooled and stained using a rat anti-mouse PE-Cy7-labelled anti-EpCAM antibody (118215, BioLegend), a rat anti-mouse FITC-labelled anti-CD45 antibody (103108, BioLegend) and DAPI (Carl Roth). Single alive EpCAM+ CD45- cells were sorted using the SH800 Cell Sorter (Sony). Isolation of barcoded RNA and subsequent cDNA library preparation were performed using the Chromium single cell 3’ reagent kits v2 and the Chromium Controller (10x Genomics), following manufacturer’s instructions. cDNA libraries were sequenced in paired end mode on a NextSeq 500 sequencer (Illumina) at the Genomics Facility, a core facility of the Interdisciplinary Center for Clinical Research (IZKF) Aachen within the Faculty of Medicine at RWTH Aachen University. Each sample corresponded to three to six animals pooled. In addition, two datasets were pooled for the analysis of the day 25 sample.

### Crypt and villi sample preparation for 3’RNA-Seq

Small intestines were rolled and snap-frozen in liquid nitrogen for 30 s to 1 min before being transferred to a -80°C freezer until further use. 12µm-thick cryosections were then cut using a Leica Cryotome and mounted onto UV-treated Zeiss PEN membrane slides. Next, slides were transferred to successive bathes of 95% ethanol, 75% ethanol, 50% ethanol, 1% cresyl violet solution in DEPC-treated water, 50% ethanol, 75% ethanol, 95% ethanol, 100% ethanol (x2) for 30 seconds each. The slides were finally left for 5 minutes in 100% ethanol before being allowed to dry for 5 minutes at room temperature. Crypts (1 million µm²) and villi (2 million µm²) were then microdissected using a Zeiss PALM MicroBeam laser microdissection system (10x objective, cut energy 57 (1-100), cut focus 58-60 (1-100)) and catapulted into opaque UV-treated Zeiss adhesive caps. RNA was immediately isolated from the microdissected tissue pieces using the microRNeasy kit (Qiagen) following manufacturer’s instructions. Finally, cDNA libraries were prepared using the QuantSeq 3’ mRNA-Seq library prep kit FWD with the i5 6nt dual indexing add-on module (Lexogen) and sequenced in single end mode on a NextSeq 500 sequencer. In addition, 4µm-thick PFA-fixed paraffin-embedded small intestinal tissue sections were stained with hematoxylin and eosin to illustrate postnatal crypt development.

### Bioinformatic analysis

RNA-Seq raw sequence reads were trimmed for adapter/low quality reads using cutadapt (version 4.4), aligned to the reference genome (mm10, GRCm38) using STAR (version 2.7.10a) and then a read count matrix was generated using the featureCounts function from the rsubreads package (version 2.14.2) ^35^. Subsequent pre-analysis and differential expression analysis of the RNA-Seq data were carried out using the DESeq2 package (version 1.38.3) ^36^. The raw read counts were normalised utilising DESeq2’s VST method. Low count genes (overall expression below 60 counts in the dataset) were filtered out and zeroes were imputed. Global and crypt-villus transcriptomic datasets were batch-corrected using ComBat-seq ^37^. To identify differentially expressed genes, a generalised linear model was employed to fit dispersion estimates, with the likelihood ratios test (LRT) used to establish statistical significance. Genes that showed an adjusted p-value (adjustment method Benjamini-Hochberg) of less than 0.05 and log_2_ fold change (FC) higher/lower than 1 were identified as differentially expressed (DE). Additionally, the ashr method was applied to shrink log_2_ fold changes ^38^.

Temporally DE genes were identified by comparing DE genes between any two consecutive timepoints (d05 vs d01; d10 vs d05; d25 vs d10). Hierarchical clustering was performed on gene expression value, the ward.D2 method was chosen empirically for gene clustering. To allow overall comparability, the fold-changes and adjusted p-values in each analysis were capped at maximum and minimum thresholds corresponding to the 0.01 and 99.9 percent quantiles (negative and positive fold-changes) and the 0.01 percentage quantile for adjusted p-values in the volcano plots. All supplementary tables contain the raw, uncapped data.

MS raw files were processed with MaxQuant software version 1.5.7.4 ^39^, peak lists were identified by searching against the mouse UniProt protein database (downloaded 2018.07.11). Searches were performed using trypsin as an enzyme, maximum 2 missed cleavages, precursor tolerance of 20 ppm in the first search used for recalibration, followed by 7 ppm for the main search and 0.5 Da for fragment ions. Carbamidomethylation of cysteine was set as a fixed modification, methionine oxidation and protein N-terminal acetylation were set as variable modifications. The required false discovery rate (FDR) was set to 1% both for peptide and protein levels and the minimum required peptide length was set to seven amino acids. Label free quantification (LFQ) based on two peptides was used. The proteomic dataset was analysed using the limma R package (version 3.54.2) ^40^. Protein intensities were log-transformed, normalised, and fitted to a linear model, with an empirical Bayes model applied to stabilise variance estimates. Differentially abundant proteins were identified using the same thresholds as described before in the RNA-Seq data analysis. To identify the origin of the proteins that were present in the proteomic dataset, but absent in the bulk RNA-Seq count matrix, we obtained the processed Seurat R-object files for mammary gland tissue published inside the *Tabula Muris* dataset ^41^. The R-object already contained annotations and clustering of cell types. This annotation was consequently used to map the uncorrelated proteins to the specific cell types of the mammary gland tissue.

For the pre-processing of the scRNA-Seq data, de-multiplexing of fastq files and alignment to the mm10 mouse genome were performed using the Cellranger toolkit (version 3.1.0) by 10x Genomics and the version 2.5.1b of STAR included in Cellranger. Quality control, normalisation, dimensionality reduction, unsupervised clustering and identification of differentially expressed genes were performed using the Seurat v4 R package ^42^. During pre-processing, cells with less than 600 genes (nFeature_RNA) and more than 25% mitochondrial counts were filtered out. Additionally, cells with fewer than 1,000 or more than 40,000 total RNA counts (nCount_RNA) were excluded to remove low-quality cells (e.g., empty droplets) and potential doublets. A subset of the 2,000 highest variable genes was identified using Seurat’s FindVariableFeatures function. Seurat’s default integration algorithm was used to combine datasets for the main analysis. Seurat processes scRNA-seq data by normalizing, identifying highly variable genes, and reducing dimensionality to cluster cells based on transcriptional similarity. Clusters were annotated based on canonical marker genes from literature (Fig. 5b). Smaller subclusters were merged to generate five clusters to accommodate to the five major small intestinal cell types (absorptive enterocytes, EECs, goblet and Paneth cells, stem cells and tuft cells). For transcription factor analysis, we used the DoRothEA R package (version 1.18.0) to estimate transcription factor (TF) activities from gene expression data. DoRothEA employs a curated collection of TF-target interactions weighted by confidence levels (A to E). TF activities were computed using the VIPER algorithm, and results were scaled using z-scores for downstream analysis. Initially, TF activity was calculated per cluster absorptive enterocytes, Goblet/Paneth, EEC and stem cells; Tuft cells were excluded due to low cell count) for each timepoint (day 1, 5, 10 and 25). To assess TF activity consistency across timepoints, we extracted significant TF activity results for the absorptive enterocyte and Goblet/Paneth clusters and identified the top 15 most active TFs per timepoint based on z-score ranking. Presence-absence matrices were generated, with presence coded as 1 and absence as 0. Heatmaps were created using the ComplexHeatmap package (version 2.22.0), and hierarchical clustering was applied to visualize TF presence patterns across timepoints ^43,44^. Three of the five identified clusters of cells (all but stem and tuft cells) were further divided into subclusters and reanalysed to explore gene expression differences within each cell population. The Harmony algorithm was used to integrate the data of the EEC subset.

^45^ This reanalysis was followed by a new marker gene analysis to identify marker genes for each subcluster. The expression of the top 50 DE genes (based on log2 fold change) of pre-weaning crypts against pre-weaning villi from our 3’ bulk RNA-Seq of crypts and villi was aggregated and overlayed onto the single cell absorptive enterocyte and Goblet/Paneth cluster to visualise potential crypt-to-villus gradients. In addition, goblet and Paneth cells were identified using the expression of *Zg16* and *Lyz1*, respectively. The EEC subclusters were annotated using known marker genes from the literature^2^. For the absorptive enterocyte and Goblet/Paneth clusters, differentially expressed marker genes (DE genes) were identified using a Wilcoxon signed-rank test with false-discovery-rate correction (Seurat’s FindMarkers method). To reduce overall complexity, only consecutive timepoints (day 5 vs day 1, day 10 vs day 5 and day 25 vs day 10) were subsequently chosen for further analysis. Overall, the same thresholds as the ones described in the RNA-Seq analysis were chosen for this DE analysis.

DE gene-based gene ontology (GO) analysis was calculated in the same manner for all datasets. A comparative cluster analysis was done using the clusterprofiler package (version 4.12.6) ^46^. Only the biological process (bp) ontology was used. Only the top significant GO terms (adj. p < 0.05 (adjustment method Bonferoni-Holm) were reported.

Pathway activities were inferred using the PROGENy (version 3.19) and decoupleR (version 2.10) R packages for the kinetic bulk RNA-Seq and proteomic datasets, ^47,48^. PROGENy computes scores for 14 signalling pathways based on gene expression data. Default settings were used, and pathway scores were scaled for comparisons. Boxplots were generated on these inferred pathway activities per timepoint, followed by an analysis of variance and a Tukey-HSD post-hoc-test.

### Sex- and gender-based analyses

Since the focus of this study was on neonatal and infant samples collected prior to puberty of the animals we did not analyse sex-specific characteristics. Both male and female neonatal and infant animals were included in the study; sex-specific genes such as *Xist*, *Jpx*, *Ftx*, *Tsx* and *Cnbp2* were removed from the datasets.

## Results

### Developmental rather than microbial factors drive global transcriptome changes during the postnatal period

First, we performed bulk RNA-Seq on total small intestinal epithelial cells isolated from 1-, 5-, 10- and 25-day-old C57BL/6J SPF mice to identify the major transcriptional changes occurring during the early postnatal life. Principal component analysis (PCA) showed a pronounced difference between the pre-weaning (1-, 5- and 10-day-old) and post-weaning (25-day-old) samples along the first dimension (PC1, 72.7%), as well as a gradual shift with increasing age among the pre-weaning samples along the second dimension (PC2, 12.3%) (Fig. 1b). Consistent with the postnatal emergence of Paneth cells and the dietary transition from breast milk to solid food, genes were most frequently differentially expressed (DE) between pre-weaning and post-weaning samples (Fig. S1a-b, Suppl. Table 1) ^30^. Whereas the increased expression in Paneth cell-associated antimicrobial genes (e.g. defensins (*Defa*), phospholipase A_2_ (*Pla2g2a*), lysozyme (*Lyz1*), regenerating family member 3γ (*Reg3g*)) occurred steadily throughout the postnatal period, upregulation of digestive enzymes (e.g. trehalase (*Treh*), sucrase isomaltase (*Sis*)) was most pronounced at weaning (Fig. 1c, Fig. S1d, Suppl. Table 1). In contrast, transmembrane transporters for iron (e.g. *Trf* and *Meltf*), folate (e.g. *Folr1*), vitamin A (e.g. *Ttr*), bile acids (e.g. *Abcb11*), and lipids (e.g. *Lrp2*) as well as glutathione peroxidases (e.g. *Gpx3*) were downregulated at weaning, in line with the particularly high metabolic demand for essential substrates as well as the adaptation to the lipid-enriched milk diet of the newborn (Fig. 1c and Fig. S1a-b, Suppl. Table 1) ^25,49–51^. Similarly, genes associated with tissue remodelling (GO terms “external encapsulating structure” (GO:0045229), “extracellular structure organization” (GO:0043062), and “extracellular matrix organization” (GO:0030198)) were downregulated from birth on (Fig. 1c, Fig. S1a-c, Suppl. Table 1). In addition, gene expression of the endocytosis adapter disabled homolog 2 (*Dab2*), the endosomal protein endotubin (*Mamdc4*) and enzymes such as cathepsin L (*Ctsl*) was at its highest during the pre-weaning period, in line with the presence of foetal-type enterocytes with intraepithelial protein digestion in supranuclear vacuoles during early life (Fig. 1c, Fig. S1a-b, Suppl. Table 1). Finally, genes downregulated between postnatal day 1 and day 5 included known foetal-specific genes (e.g. *H19*) (Fig. 1c and Fig. S1a). Fewer genes were differentially expressed between day 5 and day 10 indicating a relatively stable situation of the epithelium between the two major transcriptional shifts directly after birth and at weaning (Fig. 1c).

Pathway analysis using PROGENy revealed a transient increase in Jak/Stat signalling with a maximum at postnatal day 10, possibly related to the transient increase in mucosal cytokine expression during the so-called weaning reaction (Fig. 1d) ^15,16^. This analysis also identified an unexpected increase of the mitogen-activated protein kinase (MAPK), epidermal growth factor receptor (EGFR), and phosphoinositide 3-kinase (PI3K) pathways after weaning, whereas the NF-κB and TNF signalling pathways remained unaltered during the postnatal period (Fig. 1d and Fig. S1e). The decrease in the transforming growth factor (TGF)β after weaning is most likely due to the loss of breast milk-derived TGF-β ^52^. Interestingly, the epithelial WNT pathway decreased over time despite the delayed appearance of WNT ligand-producing Paneth cells, suggesting alternative dominant sources ^53,54^. Finally, the fact that the hypoxia-mediated signalling pathway decreased after birth before increasing again at weaning was possibly related to the delayed luminal colonization by anaerobic bacteria (Fig. 1d).

As transcriptional changes occurred for specific groups of genes at defined time points after birth (Fig. 1e), a time pattern analysis of the gene expression between consecutive time points after birth (upregulated (U) and downregulated (D) DE genes or non-regulated genes (N) between 1- and 5, 5- and 10- and 10- and 25-day-old) was performed. It confirmed the previously reported dominant transcriptional shift occurring at weaning with NND and NNU being the two largest temporal patterns, comprising 1568 and 1415 genes, respectively (Fig. 1f, Suppl. Table 2). Genes within the NND and NNU, as well as the DNU, UND and UNU temporal patterns significantly correlated with a set of genes previously reported to be differentially regulated by BLIMP1 (encoded by *Prdm1*) and MAFB, two developmentally-regulated transcription factors known to drive the intestinal epithelial neonatal-adult transition ^6–9^ (Fig. 1f). Consistent with the strong influence of developmental factors, comparative analysis of the small intestinal epithelium of age-matched pre-weaned SPF and GF mice revealed relatively minor differences with few differentially expressed genes (Fig. S1f-g, Suppl. Table 3). This was unexpected and is in marked contrast to the situation in the adult individual, where the microbiota has been identified as influencing epithelial metabolic enzyme, glycosylation and antimicrobial effector gene expression ^14, 55^.

Gene ontology (GO) analysis mainly assigned the NNU and NND time patterns to metabolic and nutrient transport processes, and to the loss of foetal-type enterocytes at weaning (Fig. 1g). In contrast, genes differentially regulated between 1 and 5 days after birth (DNN, 642 genes and UNN, 48 genes), were associated with Wnt signalling (GO:0016055), extracellular matrix organisation (GO:0030198) and tissue remodelling (GO:0043062) (Fig. 1g, suppl. Table 2). Other temporal patterns identified smaller groups with less than 100 genes each that warrant future analysis (Fig. 1f). Together, these results define the overall time course and relative contribution of epithelial transcriptional changes that accompany the establishment of mucosal host-microbial homeostasis.

### Maternally acquired proteins in the pre-weaning epithelium revealed by proteome analysis

The proteome analysis of total isolated small intestinal epithelium was based on 4562 detected proteins, of which 4160 were as well detected at the transcriptional level by bulk RNA-Seq (Fig. 2a-b). Mapping of the global small intestinal epithelial proteome of 1-, 5-, 10- and 25-day-old mice confirmed the strong difference between pre- and post-weaning samples (PC1, 47.5%; Fig. 2c). Consistently, DE protein analysis showed increased expression of Paneth cell-associated antimicrobial effector molecules (e.g. DEFA, LYZ1), metabolic (e.g. SIS, TREH, dipeptidase 1 (DPEP1), maltase glucoamylase (MGAM) and xenometabolic enzymes (e.g. carboxylesterase CES1F and CES2C), transport molecules (e.g. solute carrier SLC5A45), mitochondrial proteins (e.g. arginase 2 (ARG2), beta-carotene oxygenase (BCO2), sulfitoxidase (SUOX)), and cell proliferation (e.g. MKI67) and tight junction (e.g. claudin CLDN15) proteins at weaning (Fig. S2a-b, Suppl. Table 4). Proteins whose expression decreased at weaning were as well consistent with the previously observed transcriptional changes. Many were metabolic enzymes (e.g. argininosuccinate synthase 1 (ASS1), hexose-6-phosphate dehydrogenase/glucose-1-dehydrogenase (H6PD), N-acetylneuraminate pyruvate lyase (NPL) and GO terms catabolic/metabolic processes) or represented foetal-type enterocytes (e.g. MAMDC4, CTSL, alpha-N-acetylgalactosaminidase (NAGA) and GO terms vesicle/vacuole organisation) (Fig. 2d-e, Suppl. Table 4). PROGENy analysis of the proteomic dataset confirmed many of the changes of pathway activity observed using transcriptomics such as the JAK-STAT, MAPK, PI3K pathway (Fig. S2c).

**Figure 2:**
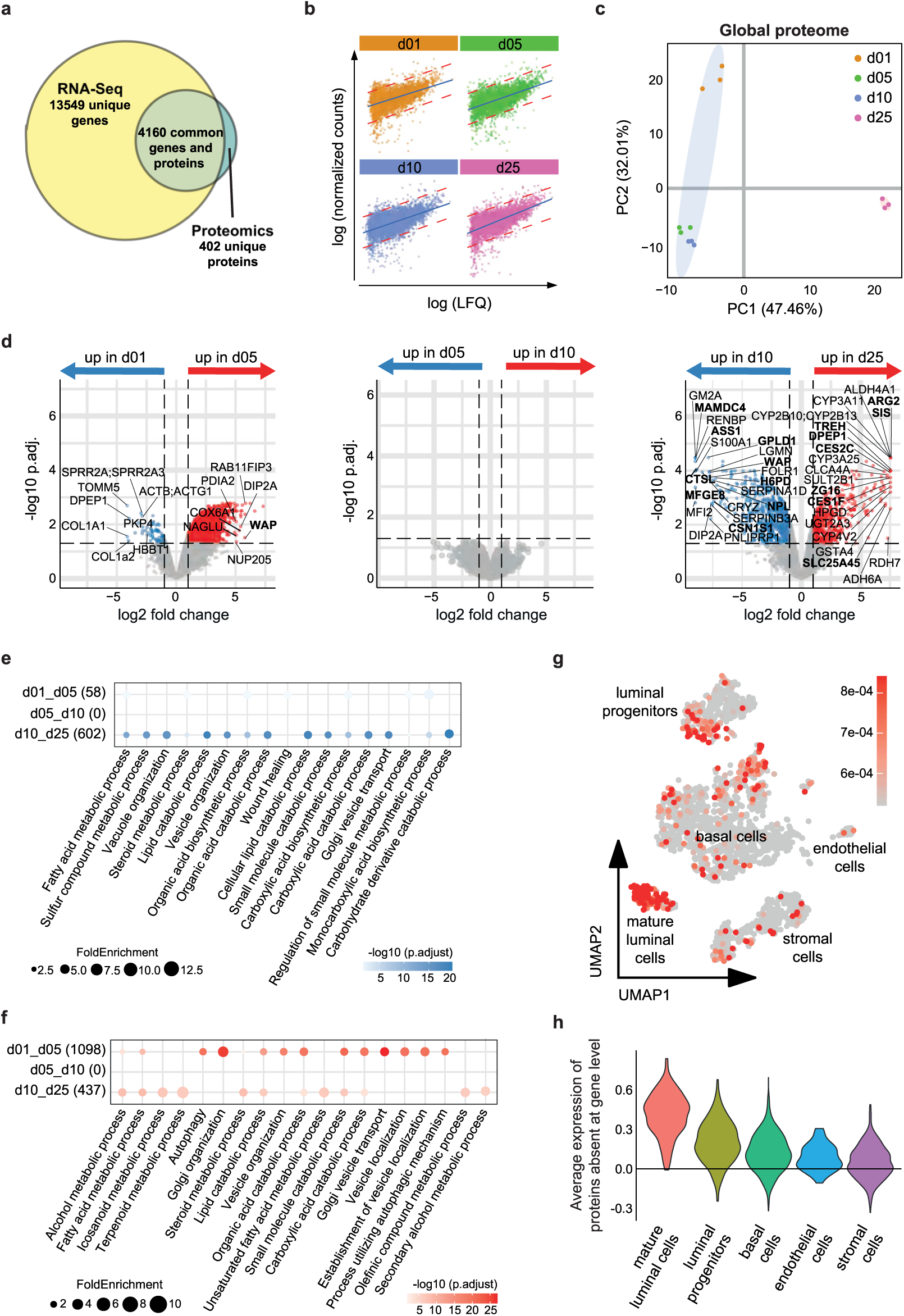
The global intestinal epithelial proteome during postnatal development. **(a)** Venn diagram representing the overlap between the RNA-Seq and proteomic datasets. **(b)** Correlation of the expression level of the 4160 common genes and proteins detected by RNA-Seq and proteomics in 1- day-old (orange), 5-day-old (green), 10-day-old (blue) and 25-day-old (pink) animals.**(c)** PCA plot representing the full small intestinal epithelial proteome of 1-day-old (n=3), 5-day-old (n=3), 10-day-old (n=3) and 25-day-old (n=3) mice. Ellipses were estimated using the Khachiyan algorithm and show the pre-weaning (light blue) and post-weaning (light pink) samples. Colour code as in Fig. 2b. **(d)** Volcano plots showing the differential expression of proteins between 1- and 5-day-old (top), 5- and 10-day-old (middle) and 10- and 25-day-old animals (bottom). Blue and red dots indicate proteins statistically (p_adj_<0.05) upregulated (|log2FC|>1) in the younger and older animals, respectively. Grey dots indicate non-significantly and/or non-differentially regulated proteins. **(e)** Dot plot showing the top 10 enriched GO terms (by adjusted p-value) for the differentially expressed genes downregulated with age. The number of DE genes associated with a GO term for each comparison is indicated in brackets. **(f)** Dot plot showing the top 10 enriched GO terms (by adjusted p-value) for the differentially expressed genes upregulated with age. The number of DE genes associated with a GO term for each comparison is indicated in brackets. **(g)** UMAP plot displaying the different types of cells in breast tissue identified by the previously published *Tabula Muris* and overlapped with the expression level of the intestinal epithelial proteins that were not detected at the transcriptional level by RNA-Seq. **(h)** Violin plot showing the quantification data of Fig. 2g. Dataset available under the identifier PXD059639 at the ProteomeXchange Consortium.

Whereas day 5 and 10 samples clustered closely together on the PCA (no DE proteins between them), they both clustered away from day 1 samples (PC2, 32.0%; Fig. 2c-d). Interestingly, this was due to a high number of proteins being upregulated (1098) rather than downregulated (58) between day 1 and day 5 (Fig. 2d, Suppl. Table 4). Most of the early upregulated proteins were found within UNN and UND, two of the main time patterns identified by the proteomic analysis (Fig. 2d, Fig. S2d-e, Suppl. Table 5). This sharply contrasted with the transcriptomic dataset which had fewer genes (126) differentially upregulated between day 1 and day 5 after birth (Fig. 1c and f). A possible explanation for this finding is the presence of milk-derived proteins (e.g. whey acid protein (WAP)) in the enterocytes of 5-day-old but not 1-day-old mice (Fig. 2d). In fact, using a previously published single cell dataset of mammary gland tissue from the *Tabula Muris*, 135 of the 402 molecules detected at the protein but not mRNA level could be matched to genes transcribed in progenitor and mature luminal cells of breast tissue, suggesting their breast milk origin (Fig. 2g-h) ^41^. Consistently, typical milk proteins (e.g. WAP, milk fat globule EGF and factor V/VIII domain containing (MFGE8), casein alpha S1 (CSN1S1)) were detected in day 10 but not day 25 samples (Fig. 2d). Another explanation for the higher number of proteins upregulated between day 1 and day 5 might be due to non-transcriptionally regulated processes like autophagy (e.g. autophagy-related 3 (ATG3), GRIP and coiled-coil domain containing 2 (GCC2), dynactin subunit 1 (DCTN1), kinesin family member 3B (KIF3B), cullin 3 (CUL3), huntingtin (HTT), GO terms autophagy (GO:0061919, GO:0006914) and vesicle localization, organization and transport (GO:0016050, GO:0048193)) and lipid catabolism and metabolism (e.g. lipase E (LIPE), fatty acid transport protein 2 (SLC27A2), caseinolytic mitochondrial matrix peptidase chaperone subunit X (CLPX), galactosidase α (GLA), arylsulfatase A (ARSA), hexokinase 2 (HK2), aminoacylase 1 (ACY1) and aspartylglucosaminidase (AGA) and GO:0016042, GO:0033559, GO:0008202) representing processes involved in the enteral intraepithelial trafficking and processing of dietary metabolites by foetal-type enterocytes (Fig. 2f, Fig. S2f and Suppl. Table 4 and 5) ^8,25^. Alternatively, the increase in protein expression may reflect the increased mTOR- and eIF2a kinase-mediated autophagosome activity induced by metabolic starvation following the interruption of maternal transplacental nutrient supply at birth ^56,57^.

### Age-dependent compartmentalisation of gene expression along the crypt-villus axis

Analysis of the global intestinal epithelium only incompletely reflects the distinct anatomical compartmentalisation and functional specialisation of the small intestinal epithelium. To account for differences in the epithelial transcriptome along the crypt-villus axis, we next combined laser capture microdissection (LMD) with bulk RNA-Seq to individually analyse the transcriptome of the crypt (or intervillus area for pre-weaning animals) and villus epithelium (Fig. 3a). Of note, although LMD was used to obtain epithelial cell material, the samples also contained transcripts from stromal (e.g. endothelin (*Edn3*)) and/or immune (e.g. immunoglobulin κ light chain (*Igkc*) and α heavy chain (*Igha*)) cells located in the vicinity of the intestinal epithelium (Fig. 3c, Fig. S3a-c).

**Figure 3:**
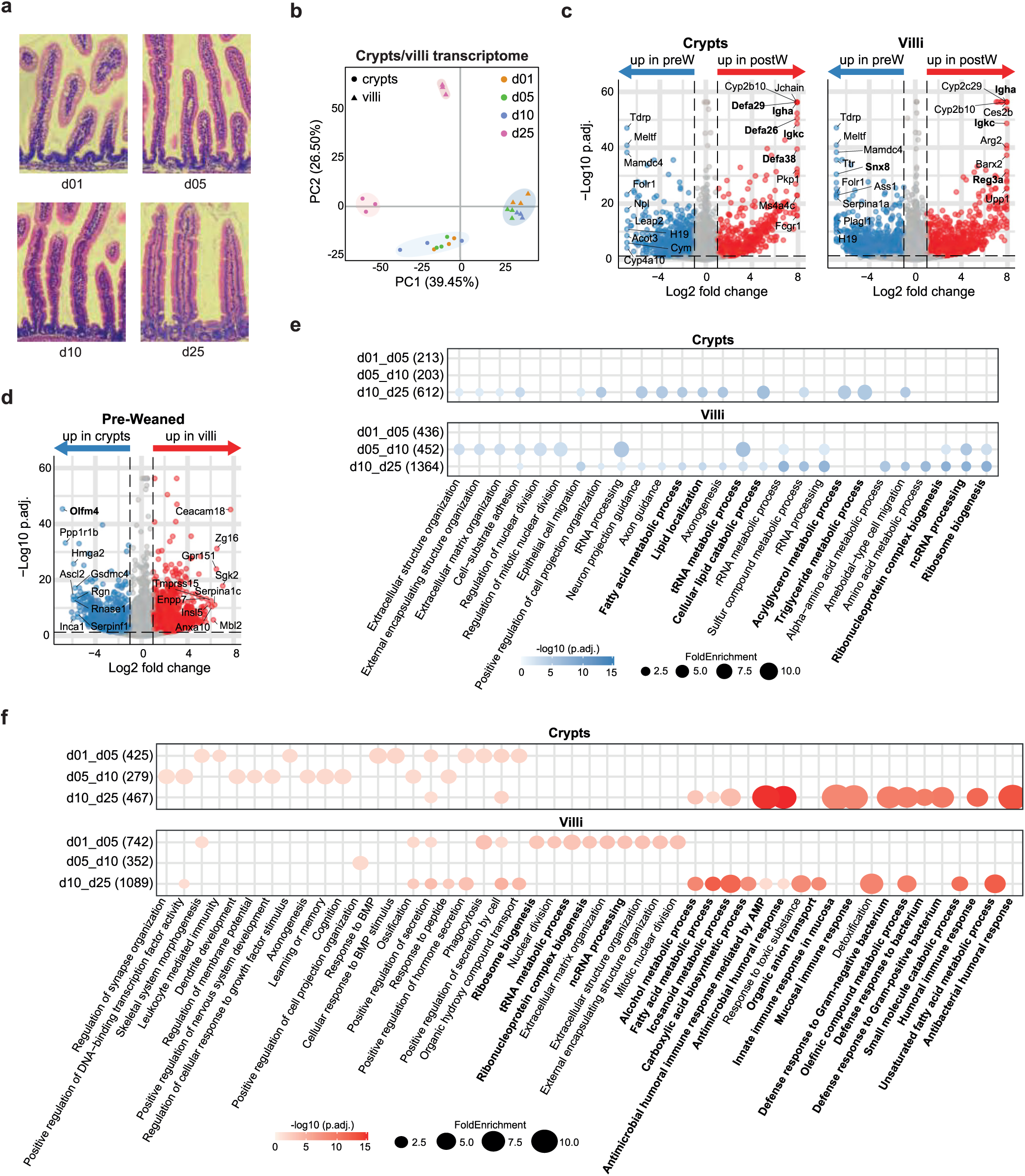
Spatial epithelial gene expression along the crypt-villus axis. **(a)** Pictures of small intestinal tissue sections obtained from 1-, 5-, 10- and 25-day-old animals and stained using H&E staining. Note the development of the intervillus/crypt structure with age. **(b)** PCA plot representing the small intestinal epithelial transcriptome of either the crypt/intervillus area (circles) or villi (triangles) of 1- day-old (orange, n=3), 5-day-old (green, n=3), 10-day-old (blue, n=3) and 25-day-old (pink, n=3) mice. Ellipses were estimated using the Khachiyan algorithm and show the pre-weaning (crypts: light blue; villi: grey blue) and post-weaning (crypts: light pink; villi: grey pink) samples. **(c)** Volcano plots showing differential gene expression between pre-weaning (1-, 5- and 10-day-old) and post-weaning (d25) crypt samples (left) and pre-weaning (1-, 5- and 10-day-old) and post-weaning (d25) villi samples (right). Blue and red dots indicate genes statistically (p_adj_<0.05) upregulated (|log2FC| > 1) in the younger and older animals, respectively. Grey dots indicate non-significantly and/or non-differentially regulated genes. **(d)** Volcano plots showing differential gene expression between pre-weaning (1-, 5- and 10-day-old) crypt and villus samples. Blue and red dots indicate genes statistically (p_adj_<0.05) upregulated (|log2FC| > 1) in the crypt and villi, respectively. Grey dots indicate non-significantly and/or non-differentially regulated genes. **(e)** Dot plot showing the top 10 enriched GO terms (by adjusted p-value) for the differentially expressed genes downregulated with age in crypts (top) and in villi (bottom). The number of DE genes associated with a GO term for each comparison is indicated in brackets. **(f)** Dot plot showing the top 10 enriched GO terms (by adjusted p-value) for the differentially expressed genes upregulated with age in crypts (top) and in villi (bottom). The number of DE genes associated with a GO term for each comparison is indicated in brackets. Dataset available under GSE283142.

PCA revealed a clear separation between the crypt/intervillus and villus epithelial transcriptomes at all ages (Fig. 3b). Irrespective of the age of the animal, genes differentially upregulated in the crypt/intervillus *versus* villus epithelium included stem cell markers (e.g. olfactomedin 4 (*Olfm4*)), molecules involved in DNA replication (e.g. PCNA clamp associated factor (*Pclaf*), topoisomerase (*Top2a*)) and antimicrobial host defence (e.g. *Defa*, deleted in malignant brain tumours 1 (*Dmbt1*)) genes (Fig. 3d, Fig. S3a and d, Suppl. Table 6).

PCA also confirmed the high similarity between the day 1, 5, and 10 samples and the strong difference between the pre- and post-weaning transcriptomes. Compared to the pre-weaning intervillus epithelium, the post-weaning crypt epithelium was characterised by GO terms associated with antimicrobial responses and represented by increased Paneth cell-associated genes (e.g. *Pla2g2a*, *Defa, Reg3b, Reg3g*), consistent with the emergence of Paneth cells during the second week of life (Fig. 3c and f, Fig. S3a-b, Suppl. Table 6) ^22,30^. Simultaneously, a decreased expression of genes represented by GO terms related to lipid metabolism was also observed, most likely representing the transition from the lipid-rich breast milk diet to solid food (Fig. 3c and e, Fig. S3a-b, Suppl. Table 6) ^58^. Additionally, the formation of deeper crypts during the late postnatal period may affect the metabolic and transport capabilities of the post-weaning crypt epithelium, due to reduced contact with dietary substrates.

In contrast, the post-weaning villus epithelium was characterised by an increased expression of genes represented by GO terms involved in nutrient digestion and transport (e.g. *Treh*, *Mgam*, nucleoside transporter *Slc28a1*, pyrimidine-degrading beta-ureidopropionase *Upb1*, cytochrome P450 *Cyp1a1* and *Cyp4b1*, monoamine oxidase A (*Maoa*)), and mucosal barrier molecules (e.g. *Reg3a*, *Muc3/Muc17)*, compared to the pre-weaning villus epithelium (Fig. 3c and f and Fig. S3a and c, Suppl. Table 6) ^59,60^. Interestingly, translational activity (GO terms “ribosome biogenesis” (GO:0042254), “tRNA metabolic process” (GO:0006399), “ncRNA processing” (GO:0034770) and “ribonucleoprotein complex biogenesis” (GO:0022613)) was first upregulated in the villus samples between day 1 and day 5, before being downregulated after weaning, possibly reflecting the increased cellular turnover and reduced need for translational activity at the villus tip upon initiation of continuous crypt-villus cell migration at weaning (Fig. 3d-f).

Furthermore, both the neonatal intervillus and villus epithelium exhibited strong signs of intracellular vesicle trafficking and enzymatic activity, as evidenced by the increased gene expression of the small GTPase *Rab30*, sorting nexin 8 (*Snx8)*, cathepsins L and B (*Ctsl* and *Ctsb* (only in the villus for the latter)), and disabled adaptor protein 2 (*Dab2*)), in line with the macromolecule internalisation and intraepithelial degradation of foetal-like enterocytes (Fig. 3c, Fig. S3a-c, Suppl. Table 6) ^8,25^. In contrast, increased expression of the IgA transport molecule polymeric immunoglobulin receptor (*pIgR*) in both post-weaning crypt and villus epithelium was consistent with the increase of IgA-secreting plasma cells occurring at weaning (Fig. 3c, Fig. S3b-c, Suppl. Table 6) ^61^.

### Transcriptional differences along the proximal-to-distal length of the small intestine

In addition to the crypt-villus axis, the small intestinal epithelium is characterised by strong anatomical and functional compartmentalisation along its length. Therefore, we additionally analysed the epithelium isolated from proximal, medial, and distal segments of the small intestine, representing the duodenum, jejunum, and ileum, respectively (Fig. 4a). The segments were well separated and anatomically defined on the PCA along the PC2 axis (16.4 %), but these segmental differences were outweighed by the previously described pre- to post-weaning transition (PC1, 64.1 %) (Fig. 4b, Fig. S4a). Hierarchical clustering of all DE genes identified 5 major clusters with differential segmental and/or temporal expression pattern (Fig. 4c). Cluster 1, 3 and 5 showed the strongest segmental differences. Thus, cluster 1, characterised by GO terms “sister chromatid segregation” (GO:0000070), “nuclear division” (GO:0000280) and “chromosome segregation” (GO:0007059), was represented by genes with a higher expression in all segments after weaning, but already in the proximal part at day 10 after birth, indicating that stem cell and TA cell proliferation started earlier in the proximal intestine (Fig. 4c-d, Fig. S4b-c, Fig. S4e, Suppl. Table 7). Cluster 3 encompassed genes whose expression peaked at birth (1-day-old), more strongly and consistently in the proximal part, and that were characterised by GO terms such as extracellular structure/extracellular matrix organisation (GO:0043062 and GO:0030198) and digestive tract/system development (GO:0048565 and GO:0055123), indicating early tissue remodelling and maturation, a prerequisite for crypt formation (Fig 4c-d, Fig. S4b-d, Suppl. Table 7). Genes of cluster 5, represented by GO terms such as “lytic vacuole organization” (GO:0080171) or “lysosome organization” (GO:0007040) most likely representing foetal-type enterocytes, were more prominently and consistently expressed in the pre-weaning distal intestine, in line with the persistence of these enterocytes in the distal small intestine during the first two weeks of life (Fig. 4c-d, Fig. S4b-d, Suppl. Table 7) ^25^. Cluster 2, characterised by GO terms associated with antimicrobial response (GO:0019730) and mucosal immune defence/defence response to bacteria (GO:0002385 and GO:0042742) and most likely representing Paneth cell-derived antimicrobial molecules, was comprised of genes uniformly upregulated in all segments post-weaning (Fig 4c-d, Fig. S4c, Fig. S4e). In contrast, genes from cluster 4 were upregulated in all segments during pre-weaning and were mostly associated with metabolic processes (Fig. 4c-d, Fig. S4c-d, Suppl. Table 7).

**Figure 4:**
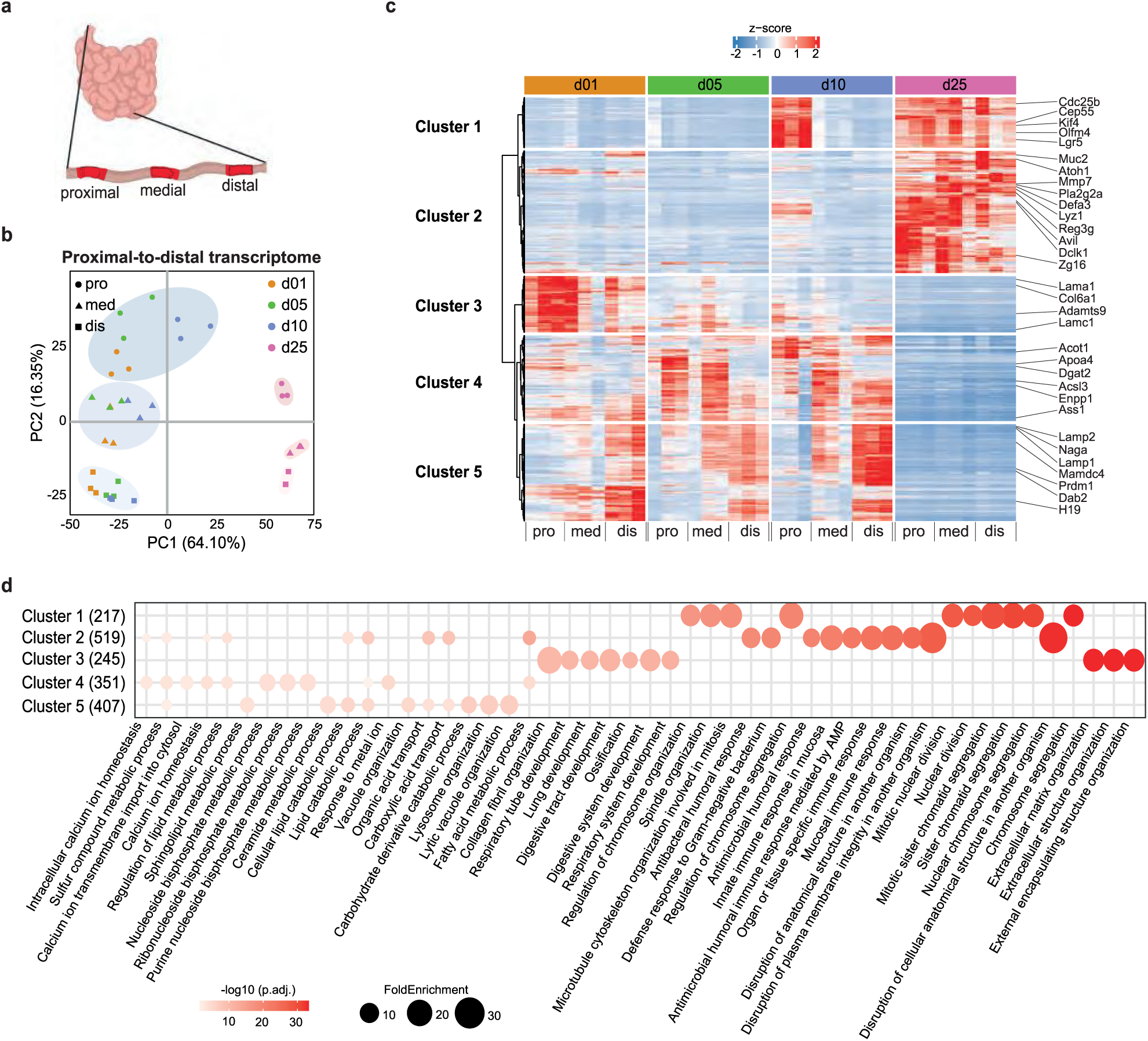
Spatial epithelial gene expression along the length of the small intestine. **(a)** Schematic representation of the small intestine showing the different samples collected. Figure created using Biorender.com. **(b)** PCA plot representing the epithelial transcriptome of either the proximal (circles), medial (triangles) or distal (squares) part of the small intestine of 1-day-old (orange, n=3), 5-day-old (green, n=3), 10-day-old (blue, n=3) and 25-day-old (pink, n=3) mice. Ellipses were estimated using the Khachiyan algorithm and show the distribution of the pre-weaning (proximal: light blue; medial: blue; distal: grey blue) and post-weaning (proximal: light pink; medial: pink; distal: grey pink) samples. **(c)** Heatmap depicting the expression level of all genes differentially regulated between the proximal and medial part or the medial and distal part of 1-, 5-, 10- and/or 25-day-old animals. **(d)** Dot plot showing the top 10 enriched GO terms (by adjusted p-value) for the differentially expressed genes associated with each of the 5 clusters identified in Fig. 4c. The number of DE genes associated with a GO term for each comparison is indicated in brackets. Dataset available under GSE283143.

**Figure 5:**
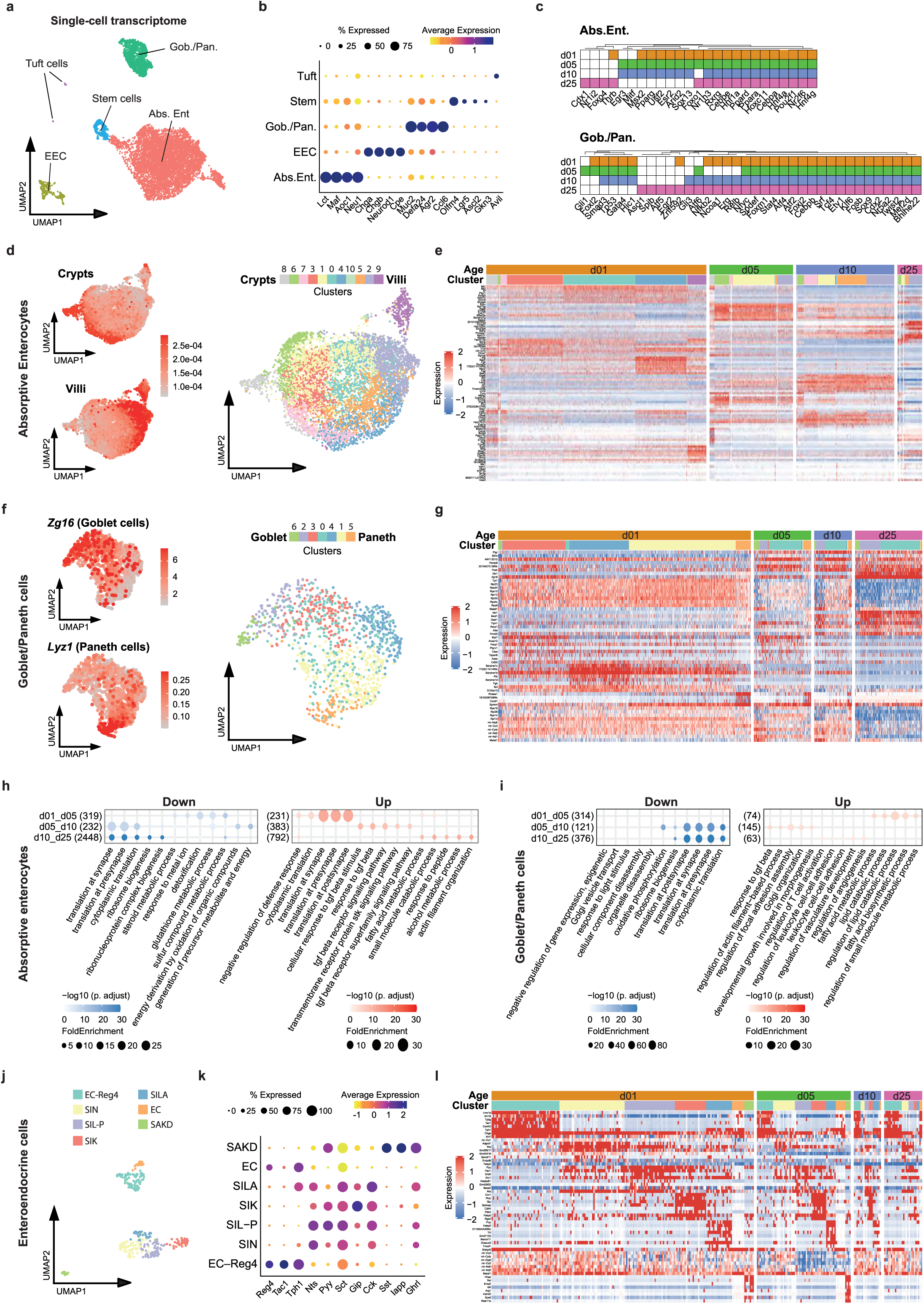
Single cell transcriptional analysis of intestinal epithelial cells. **(a)** UMAP plot displaying the different types of epithelial cells recovered from uninfected pre- and post-weaning murine small intestines: Absorptive enterocytes (red), Goblet/Paneth (green), stem (blue), enteroendocrine (ochre) and tuft (purple). **(b)** Dot plot displaying the marker genes for each cluster identified in Fig. 5a. **(c)** Graphical representation of the top 15 transcription factors (TF) identified by DoRothEA for 1- (orange), 5- (green), 10- (blue) and 25-day-old (pink) absorptive enterocytes (top) and Goblet/Paneth cells (bottom). Coloured cells depict transcription factors predicted by DoRothEA for each age group. White cells represent TF that were not detected by DoRothEA. **(d)** UMAP plots showing the expression level of genes previously found to be associated with pre-weaned crypts (top left) and pre-weaned villi (bottom left), and the different subclusters recovered from the absorptive enterocyte cluster (right). The red colour intensity of the left plots reflects the overall gene expression level. **(e)** Heatmap representing the top 8 DE genes per absorptive enterocyte subcluster. Age/cluster colour code as in Fig. 5c/Fig. 5d. **(f)** UMAP plots showing the expression level of known goblet (*Zg16*, top left) and Paneth cell (*Lyz1*, bottom left) markers, and the different subclusters recovered from the Goblet/Paneth cluster (right). The red colour intensity of the left plots reflects the gene expression level. **(g)** Heatmap representing the top 8 DE genes per Goblet/Paneth subcluster. Age/subcluster colour code as in Fig. 5c/Fig. 5f. **(h-i)** Dot plot showing the top 5 enriched GO terms (by p._adj._) for the genes differentially downregulated (left) and upregulated (right) with age in absorptive enterocytes (h) and in Goblet/Paneth cells (i). The number of DE genes associated with a GO term for each comparison is indicated in brackets. **(j)** UMAP plots showing the different subclusters recovered from the EEC cluster. **(k)** Dot plot displaying the marker genes for each cluster identified in Fig. 5i. **(l)** Heatmap representing the top 8 DE genes per EEC subcluster. Age/subcluster colour code as in Fig. 5c/Fig. 5j. Dataset available under GSE284074.

### Age- and cell type-specific developmental trajectories revealed by single cell transcriptional analysis

Next, we performed single-cell RNA sequencing (scRNA-seq) and analysed 9244 high quality cells isolated from 1-, 5-, 10- and 25-day-old mice, as well as from *Salmonella*-infected animals (4 days post-infection (p.i.)/5-day-old) (Fig. S5a-c). The single cell epithelial transcriptome of healthy animals captured all major intestinal epithelial cell types (i.e. absorptive enterocytes, goblet and Paneth cells, tuft cells, enteroendocrine cells, and stem cells), identified using established marker genes (Fig. 5a-b)^2^. Notably, epithelial cells from adult mice showed an overall lower viability. Isolation and flow cytometric sorting of viable intestinal epithelial cells from both crypts and villi proved challenging in post-weaning samples leading to an underrepresentation of crypt-based stem and Paneth cells in the day 25 sample (Fig. S5c). We initially focussed on the effect of age on epithelial cell type differentiation since, although studied in the adult host, little is known about regulatory circuits that drive epithelial cell differentiation during the neonatal and infant period ^2,20^. We employed DoRothEA, a curated collection of transcription factor (TF) gene interactions, to identify cell type specific TF activity ^62^. This allowed the identification of age-dependent TF activity specifically in absorptive enterocytes, goblet/Paneth cells, EECs and stem cells (Fig. S5d and Suppl. Table 8). A subsequent assembly of the TF with the strongest activity in each age group for absorptive enterocytes and goblet/Paneth cells revealed mainly three patterns (Fig. 5c and Fig. S5e): Firstly, TF with age-independent activity such as Hnf1a, Hnf4a, and Cebpe for absorptive enterocytes or Spdef, Tcf4, and Sox9 for goblet/Paneth cells.

Secondly, TF with post-weaning activity such as Cdx1 and SpiB for absorptive enterocytes and goblet/Paneth cells, respectively. And finally, TF with exclusive pre-weaning activity such as Msx2, Esr2, Arid2 and Sox13 that have not previously been associated with intestinal epithelial cell differentiation in absorptive enterocytes or Gata4 and Smad3 that have been identified, but not associated with the neonatal period in goblet/Paneth cells ^12,63^.

Next, we analysed in more detail the cell type clusters with high cell numbers by subclustering. A spatial dimension was added to the absorptive enterocyte subclusters by overlapping the gene expression signatures of crypt and villus epithelium previously identified by bulk RNA-Seq (Fig. 3) onto our single cell dataset (Fig. 5d). This allowed the identification of the gene signatures of single absorptive enterocytes along the crypt-villus axis and illustrated the continuous gradient of cell differentiation and functional diversity. Notably, cells obtained at different age contributed markedly differently to individual subclusters (Fig. S5f) and the heatmap illustrating the top 8 DE genes per subcluster highlighted major age-specific differentiation trajectories (Fig. 5e). A similar phenomenon was also observed for goblet/Paneth cells. Fully differentiated cells within this cluster were identified by the expression of *Zg16* and *Lyz1*, two established markers for goblet and Paneth cells, respectively (Fig. 5f). Cluster assignment was consistent with a crypt-restricted localisation of Paneth cells but global distribution of goblet cells (Fig. S5h). Cells obtained at different ages again contributed markedly differently to individual subclusters (Fig. S5g) and the heatmap illustration of the top 8 DE genes per subcluster demonstrated major differences, especially in the intermediate cell stages of 1-day-old as compared to older mice, suggesting persistent foetal-derived intermediate cells in 1-day-old mice (Fig. 5g). GO term analysis of the age-dependent transcriptional profiles identified the previously observed developmental changes of metabolic processes in both absorptive enterocytes and goblet/Paneth cells (e.g. GO terms “fatty acid metabolic process” (GO:0006631) or “alcohol metabolic process” (GO:0006066)) and confirmed an early postnatal upregulation in absorptive enterocytes and subsequent downregulation of translational activity in absorptive enterocytes and goblet/Paneth cells (Fig. 5h-i-; Fig. 2e-f; Fig. S1c-d; Fig. 3e-f, Suppl. Table 9). The increase in TGFβ activity in absorptive enterocytes between day 5 and 10 after birth (e.g. GO terms “response to tgf beta” (GO:0071559) or “tgf beta receptor signalling pathway” (GO: 0007179)) was not detected in the pathway analysis based on the global transcriptomic and proteomic profiles during this time window, highlighting the importance of cell type-specific analyses (Fig. 5h-i; Fig. 1d; Fig. S2c, Suppl. Table 9). In contrast to absorptive enterocytes and goblet/Paneth cells, the seven identified EEC subclusters assigned to seven known EEC subtypes based on previously reported marker genes were detected at all ages with similar abundancy (Fig. 5j-k, Fig. S5i-j)^2^. Additionally, the heatmap illustrating the top 8 DE genes per subcluster/subtype showed the typical “staircase” pattern for each age group, suggesting similar developmental trajectories in neonatal and adult animals for EECs, in contrary to absorptive enterocytes and goblet/Paneth cells (Fig. 5l).

### The global and cell type-specific gene response to enteric infection

To investigate epithelial gene expression under challenged conditions, we also analysed intestinal epithelial cells collected at day 4 p.i. from mice infected orally at day 1 after birth with 100-300 CFU *S.* Typhimurium (i.e. age-matched to the day 5 time point) ^64^. Mapping of the small intestinal epithelial bulk transcriptome of 5-day-old *S.* Typhimurium-infected mice on that of 1-, 5-, 10- and 25-day-old healthy mice illustrated the major infection-induced transcriptional changes (PC1, 43.5 %) consistent with our previous report (Fig. 6a) ^65^. DE gene analysis of day 5 uninfected and day 5 infected samples revealed upregulation of typical epithelial antimicrobial host response genes such as c-type lectins (e.g. *Reg3g*, *Reg3b*, *Reg3a*), alarmins (e.g. *S100a8*/*S100a9)*, inducible NO synthase 2 (*Nos2*), NAPDH oxidase 1 (*Nox1*), serum amyloid A1 (*Saa1*), and cytokines (e.g. *Tnf* and C-C motif chemokine ligand 8 (*Ccl8*)), but also a number of catabolic enzymes (e.g. tryptophan catalytic enzyme indoleamine 2,3- dioxygenase 1 (*Ido1*), uridine degrading enzyme uridine phosphorylase 1 (*Upp1*)) (Fig. 6b, Suppl. Table 10). UMAPs representing the scRNA-Seq of the uninfected (d05) and infected datasets (d05i) illustrated a marked decrease in the abundance of absorptive enterocytes in 5-day-old infected animals. Given the previously shown largely preserved tissue architecture in infected animals, with only low numbers of cleaved caspase 3 positive enterocytes, this suggests that the inflammatory environment *in vivo* aggravated the cell stress during cell isolation and flow cytometric sorting with subsequent cell exclusion from the analysis due to low data quality (Fig. 6c, Fig. S5c) ^64,66^. Interestingly, a reduced cell abundance was not observed for stem, Paneth/goblet, tuft or enteroendocrine cells (Fig. 6c, Fig. S5c), indicating a higher resilience of these cells towards the proapoptotic inflammatory environment *in vivo*. A subsequent cell type-specific analysis of the response to infection revealed sets of genes altered by infection in multiple cell types, but as well sets of genes unique to specific cell types (Fig. 6d and Fig. S6a-d, Suppl. Table 10). GO term analysis of these uniquely induced genes suggested that stem cells specifically responded to infection by increasing their metabolic activity (e.g. GO term “hexose” or “glucose metabolic process”, GO:0019318 or GO:0006006), whereas absorptive enterocytes were associated with a higher cell division rate (e.g. “chromosome segregation”, GO:0007059) and goblet/Paneth cells with a higher innate immune response orchestrated by pattern recognition receptors (e.g. “activation of innate immune response”, GO: 0002218). Lastly, an increased response to calcium ions, coupled with a higher translational and post-translational activity (e.g. “response to calcium ion”, GO: 0051592) suggested a greater stimulatory activity in infected enteroendocrine cells (Fig. 6e, Suppl. Table 10).

**Figure 6:**
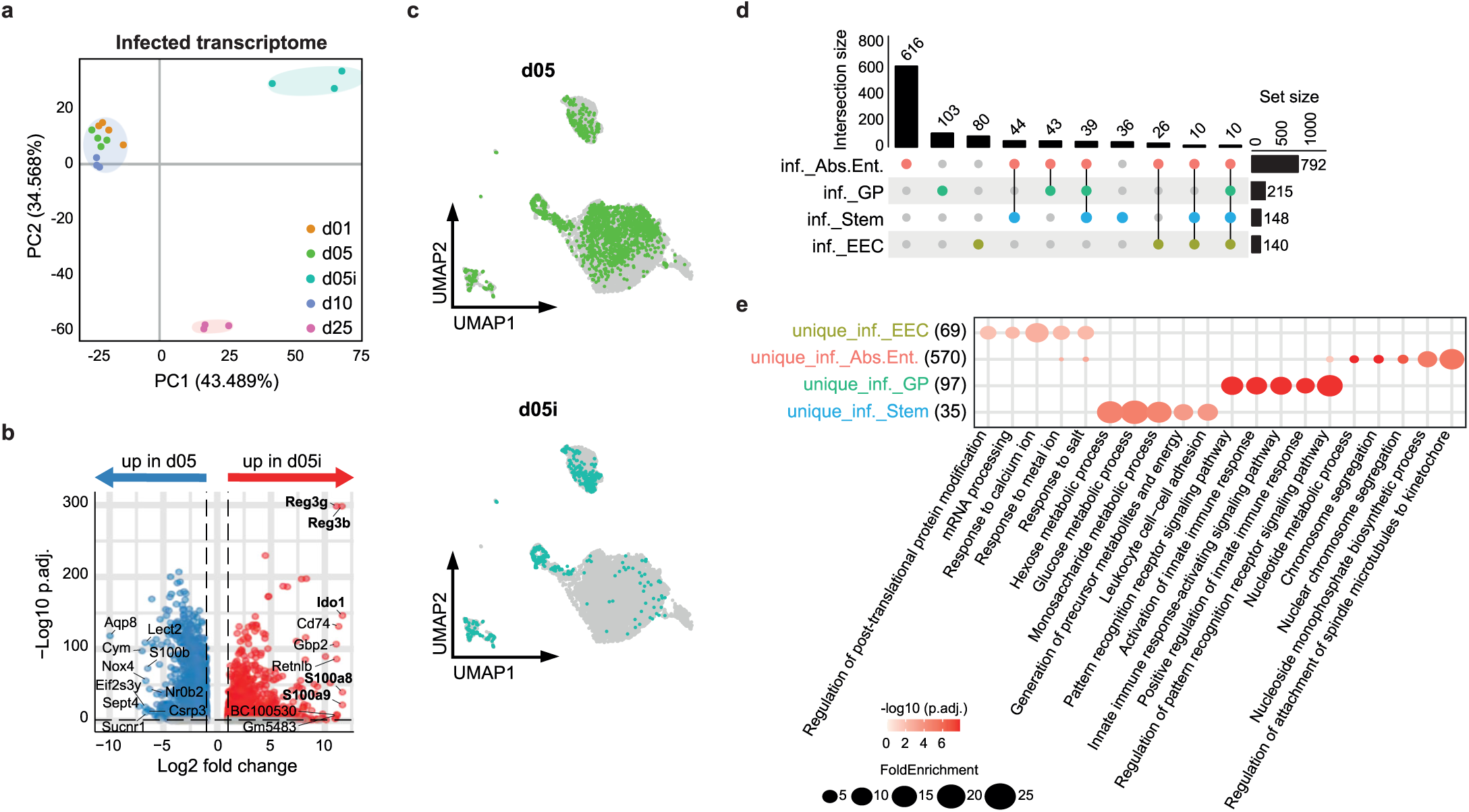
Transcriptional analysis of intestinal epithelial cells isolated from *Salmonella*-infected animals. **(a)** PCA plot representing the epithelial transcriptome of the small intestine of 1-day-old (orange, n=4), 5-day-old (green, n=4), 10-day-old (blue, n=3) and 25-day-old (pink, n=3) uninfected mice, as well as 5-day-old mice infected at birth with *S.* Typhimurium (turquoise, n=3). Ellipses were estimated using the Khachiyan algorithm and show the distribution of the pre-weaning (light blue) and post-weaning (light pink) uninfected and 5-day-old infected (light turquoise) samples. **(b)** Volcano plots showing differential gene expression between 5-day-old *S*. Typhimurium-infected and uninfected samples. Blue and red dots indicate genes statistically (p_adj_<0.05) upregulated (|log2FC| > 1) in the uninfected and infected animals, respectively. Grey dots indicate non-significantly and/or non- differentially regulated genes. **(c)** UMAP plots displaying the epithelial cells isolated from uninfected (top; green; same dataset as in Fig. S5b) and *Salmonella*-infected (bottom; turquoise) 5-day-old newborn mice. **(d)** Upset plot showing the number of genes differently regulated by one or more epithelial cell type upon infection with *S*. Typhimurium. Absorptive enterocytes (Abs. Ent.; red), goblet/Paneth cells (GP; green), stem cells (blue), enteroendocrine cells (EEC; ochre). Filled circles represent genes unique to a specific cell type, open circles represent genes shared with other cell types. Interactions with less than 10 DE genes are not shown. **(e)** Dot plot showing the top 5 enriched GO terms (by adjusted p-value) for the infection-induced DE genes that were unique to each intestinal epithelial cell type. The number of DE genes associated with a GO term for each comparison is indicated in brackets. Datasets available under GSE283495 and GSE284074.

## Discussion

Our results illustrate the complexity and striking heterogeneity of the different neonatal small intestinal epithelial cell types, demonstrate intriguing alterations in epithelial gene and protein expression early after birth and identify functional changes that may contribute to the postnatal transition. Unexpectedly, transcriptional differences between the proximal and distal intestine but also between the crypt and villus epithelium were as pronounced in the neonate as compared to the adult host despite the lack of an organ site-specific microbiota composition, the lack of defined small intestinal crypts and a reduced mucus production, making the neonatal mucosal architecture less compartmentalised ^23,64,67^. Also, whereas the enteric microbiota has a significant impact on the epithelial expression of host defence, glycosylation and metabolic genes in adult animals, our epithelial transcriptome suggested little differences between SPF and GF mice in pre-weaning animals ^14^. The low diversity and large interindividual variation of the early microbiota composition may fail to provide reliable innate immune signals, thus favouring developmental programming of the neonate intestinal epithelium ^67,68^.

Although the establishment of intestinal epithelial stem cell organoids representing all epithelial cell lineages has been a major breakthrough in intestinal epithelial cell research ^69^, organoids grown *in vitro* still lack the influence of the enteric microbiota, dietary, nervous and immune cell signals, as well as hormones and tissue ontogeny, supporting the need to analyse *ex vivo* isolated primary cells ^19,70^. Nonetheless, the isolation and analysis of primary small intestinal epithelial cells remain technically challenging. Firstly, detachment from the basal membrane, a prerequisite for the isolation and analysis of epithelial cells, rapidly reduces cell viability. While this issue is negligeable when isolating epithelial sheets for bulk RNA-Seq and proteomic analyses, it is particularly strong when isolating single cells for scRNA-Seq analysis ^26,64,66^ and this loss of cell viability during isolation may be further aggravated by tissue inflammation occurring upon infection ^71^. Secondly, the anatomical organisation of the intestinal epithelium makes it difficult to simultaneously isolate representative fractions of viable cells from villi and crypts at different ages ^72^. As our analysis focused on the early postnatal period, we prioritised villus cell viability over adult crypt cell coverage which explains the underrepresentation of crypt stem and Paneth cells in our adult sample. However, the crypt epithelium of adult mice has been previously characterised ^2,3^.

All intestinal epithelial cell types arise from Lgr5^+^ pluripotent stem cells ^10,11,73^. Notch-mediated lateral inhibition and a complex network of epithelial and mesenchymal signals and transcription factors drive differentiation into the specific cell type lineages ^1,23,74^. Many largely uncharacterised transcription factors were significantly associated with specific epithelial cell types in the neonate and await further analysis. Interestingly, foetal transcription factors were shown to reemerge and influence cell differentiation in the inflamed infant intestine ^75^. In addition, migration along the crypt-villus axis exposes cells to spatial gradients of proliferation and differentiation-regulating factors ^23^, and a proximal-to-distal gradient, while under marked transcriptional control, may also be influenced by differences in the concentration of microbial or dietary stimuli ^12,76^.

Major differences in the intestinal epithelial gene and protein expression were observed at weaning ^6–9^, largely explained by the transcriptional influence of two of the major transcriptional regulators of the pre- to post-weaning transition, BLIMP1 (*Prdm1*) and MAFB ^6–9^. They included changes in the expression of digestive enzymes reflecting the transition of breast milk to solid food ^2,7^. Cessation of breast milk feeding and thereby cessation of contact with the breast milk constituents EGF and TGFβ reduced TGFβ and EGF receptor signalling in epithelial cells after weaning ^58^. Also, the emergence of crypts and crypt-based Paneth cells was identified by the upregulation of typical Paneth cell genes such as α-defensins and phospholipase A_2_ ^22,30,73^. The transcriptional changes of both digestive enzymes and antimicrobial molecules were found throughout the intestinal tract. In contrast, epithelial cell proliferation, as a sign of the establishing crypt niche, started earlier in the proximal part of the intestine consistent with the proximal-to-distal developmental wave ^1^. The previously described “weaning reaction” in the mucosal tissue that reflects immune maturation and is accompanied by elevated expression of pro-inflammatory cytokines may be responsible for enhanced epithelial signalling via the Jak/Stat, MAPK and PI3K pathway ^15^.

Important transcriptomic and proteomic changes were also detected earlier, between day 1 and 5 after birth. In contrast to the transcriptomic analysis, the proteome analysis revealed mostly proteins upregulated between day 1 and 5 after birth. In part, this conundrum could be explained by the presence of proteins produced by mature luminal cells in the breast gland ^41^. Milk proteins in isolated intestinal epithelial cells most likely reflects the presence of foetal-type enterocytes that internalise luminal proteins by endocytosis followed by intraepithelial degradation by cathepsins ^25^. Foetal-type enterocytes were restricted to the pre-weaning period and their gene signature was most abundant in the distal part of the small intestine, consistent with the literature ^25^. In addition, the detection of proteins in the absence of the corresponding transcript may be due to increased autophagocytotic activity in the neonatal host, providing peptides amenable to mass spectrometric detection. Autophagy has been shown to be critical to overcome the energy shortage of the neonatal host imposed upon birth and until establishment of enteral feeding ^56,57^. Both aspects illustrate the unique situation of the neonatal host and identify important strategies to adapt to postnatal life.

Together, we used state-of-the-art methods and models to characterise the global, organ- and site-specific as well as single-cell transcriptional and proteomic profile of the gut epithelium at different time points after birth. Our results identify and characterise key temporal and organ site-specific gene expression profiles, determine age-dependent signalling pathway activity, and define age- and cell-type-specific transcription factor activity illustrating the strong developmental influence and only marginal effect of the enteric microbiota. We describe the global and cell-type-specific response to neonatal enteric infection and characterise the age-dependent developmental trajectories of individual cell lineages highlighting the compartmentalized and discontinuous transcriptional profiles of specific cell type. Our results provide the basis for future investigations and may help to identify strategies to reduce childhood mortality and promote mucosal homeostasis and long-term health.

## Data Sharing Plan

The bulk RNA-Seq data in Fig. 1, S1, 3, 4 and 6 and the single cell RNA-Seq data in Fig. 5 and 6 on the age, organ site, and anatomical location dependent as well as infection and microbiota-driven transcriptional changes are available in Gene Expression Omnibus NCBI (GEO) at https://www.ncbi.nlm.nih.gov/gds with the accession no. GSE283142 GSM8656671-GSM8677457), GSE283143 (GSM8656695-GSM8656730), GSE283495 (GSM8663964-GSM8663992) and GSE284074 (GSM8677452-GSM7688487). Mass spectrometry proteomics data have been deposited to the ProteomeXchange Consortium via the PRIDE ^77^ partner repository with the dataset identifier PXD059639.

## Code availability

The code used for this study is available on GitHub under https://github.com/jschoeneich/epithelial_cell_diversity. The repository includes all necessary scripts for reproducing the analyses.

## Acknowledgements

We thank Regina Holland, Martina Leufgens, Simone Martin, and Josefine Weber-Heynemann from the Institute of Medical Microbiology and Till Braunschweig, Melanie Mitchell and Ursula Schneider from the Institute of Pathology for technical help as well as Lisa Meier and Nadine Utes for excellent support with our animal husbandry. We further thank Susan A.V. Jennings and Tom Clavel for providing tissue from germ-free mice and Fabian Nienhaus for the small intestine illustration (all RWTH University Hospital Aachen). This project was supported by the core facility Genomics of the Faculty of Medicine at RWTH Aachen, Germany.

## Author contributions

I.G.C and M.W.H contributed to the conception of the work, J.S., A.D., I.G.C. and M.W.H. contributed to the design of the work, A.D., S.S., M.A.S., I.R., S.J., F.I. and C.K. contributed to the acquisition of the data, J.S., A.D., N.M.K.P.G., M.C., T.M., S.J., C.T., A-E.S., T.P., I.G.C. and M.W.H. contributed to the analysis of the data, J.S., A.D., T.P., I.G.C. and M.W.H. contributed to the interpretation of the data, J.S., A.D. and M.W.H. have drafted the manuscript, and J.S., A.D., S.S., M.A.S., T.M., C.K., T.P., I.G.C. and M.W.H. have revised the manuscript.

## Supplementary figure legends

**Supplementary Figure 1:**
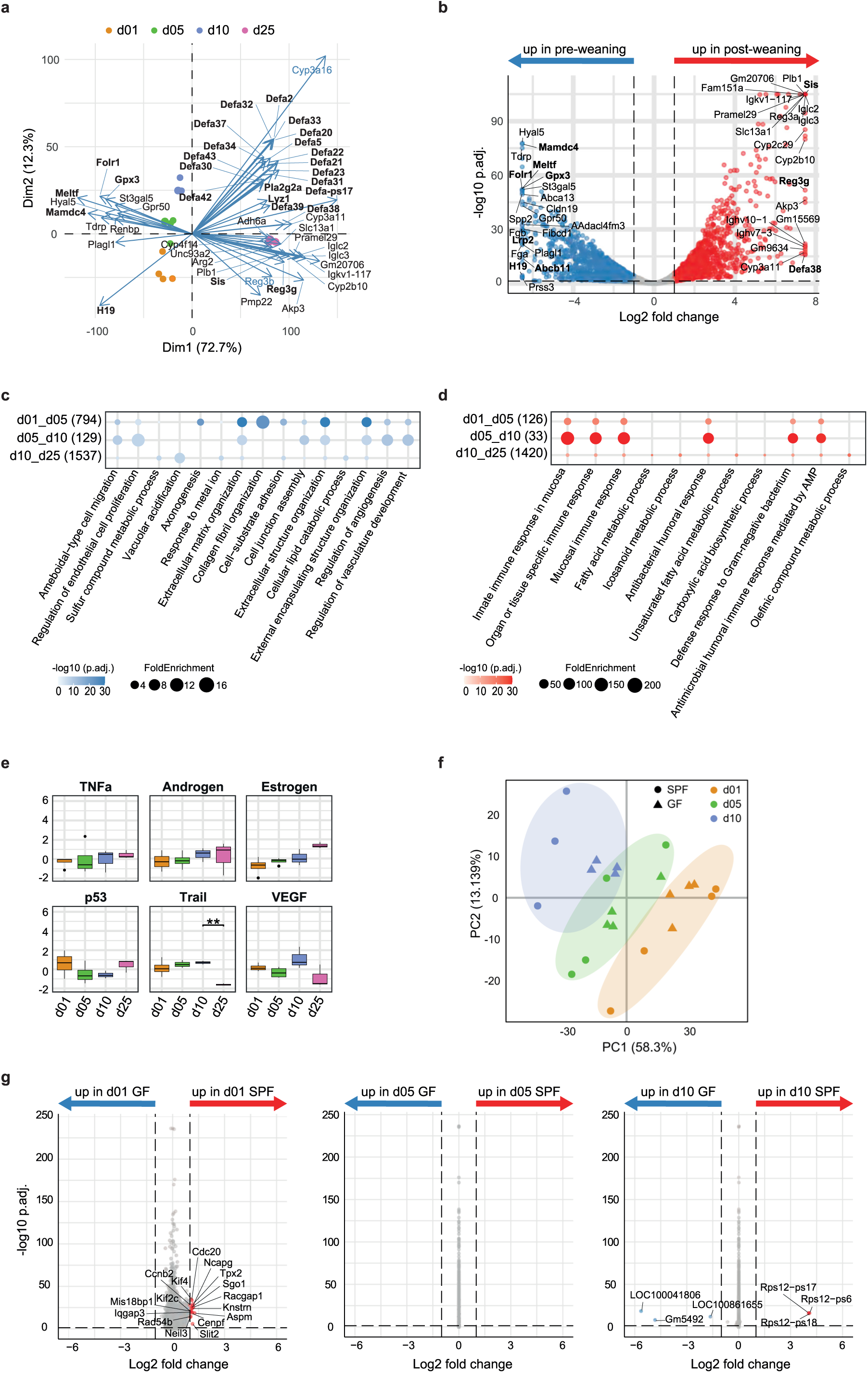
**(a)** Biplot displaying the top 50 driver genes for the 1^st^ and 2^nd^ dimension of the PCA plot shown in Fig. 1b. Black/blue names represent genes that were identified as DE/non-DE between pre- and post-weaning samples. 1-day-old (orange dots), 5-day-old (green dots), 10-day-old (blue dots) and 25-day-old (pink dots). **(b)** Volcano plots showing differential gene expression between pre-weaning (1-, 5- and 10-day-old) and post-weaning (25-day-old) samples. Blue and red dots indicate genes statistically (p_adj_ < 0.05) upregulated (|log2FC|>1) in the younger and older animals, respectively. Grey dots indicate non-significantly and/or non-differentially regulated genes. **(c)** Dot plot showing the top 5 enriched GO terms (by adjusted p-value) for the differentially expressed genes downregulated with age (blue DE genes from Fig. 1c). The number of DE genes associated with a GO term for each comparison is indicated in brackets. **(d)** Dot plot showing the top 5 enriched GO terms (by adjusted p-value) for the differentially expressed genes upregulated with age (red DE genes from Fig. 1c). The number of DE genes associated with a GO term for each comparison is indicated in brackets. **(e)** Box plots showing PROGENy pathway analysis at the different ages. Colour code as in Fig. S1a. **(f)** PCA plot representing the full small intestinal epithelial transcriptome of 1-day-old (orange, n=4/4 SPF/GF), 5- day-old (green, n=4/4) and 10-day-old (blue, n=3/4) specific pathogen free (SPF, circles) and germ-free (GF, triangles) mice. Ellipses were estimated using the Khachiyan algorithm and show the distribution of samples for each age group. **(g)** Volcano plots showing differential gene expression between GF and SPF mice at 1 (left), 5 (middle) and 10 (right) days of age. Blue and red dots indicate genes statistically (p_adj_<0.05) upregulated (|log2FC| > 1) in GF and SPF animals, respectively. Grey dots indicate non-significantly and/or non-differentially regulated genes.

**Supplementary Figure 2:**
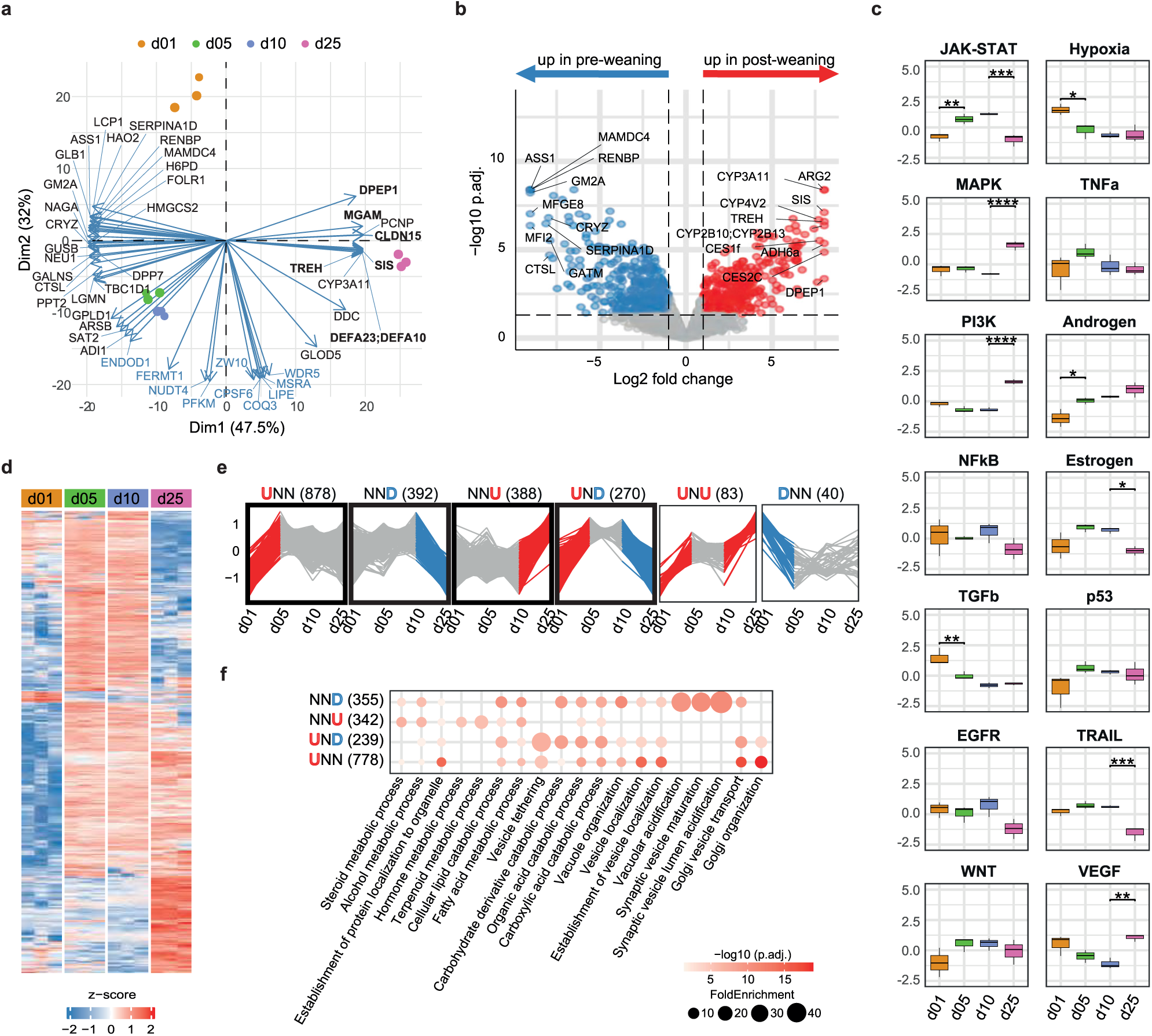
**(a)** Biplot displaying the top 50 driver proteins for the 1^st^ and 2^nd^ dimension of the PCA plot shown in Fig. 2c. Black/blue names represent proteins that were identified as DE/non-DE between pre- and post-weaning samples. 1-day-old (orange dots), 5-day-old (green dots), 10-day-old (blue dots) and 25-day-old (pink dots). **(b)** Volcano plots showing differential protein expression between pre-weaning (1-, 5- and 10-day-old) and post-weaning (25-day-old) samples. Blue and red dots indicate proteins statistically (p_adj_<0.05) upregulated (|log2FC| > 1) in the younger and older animals, respectively. Grey dots indicate non-significantly and/or non-differentially regulated proteins. **(c)** Box plots showing PROGENy pathway analysis at the different ages. Colour code as in Fig. S2a **(d)** Heatmap depicting the expression level of all proteins differentially regulated between either 1- and 5-day-old, 5- and 10-day-old and/or 10 and 25-day-old animals. **(e)** Six most abundant temporal postnatal protein expression patterns as identified by temporal DE analysis. The number of DE proteins for each pattern is indicated in brackets. **(f)** Dot plot showing the top 5 enriched GO terms (by p_adj_ value) for the differentially expressed genes identified as belonging to the NND, NNU, UND and UNN expression patterns. The number of DE proteins associated with a GO term for each pattern is indicated in brackets.

**Supplementary Figure 3:**
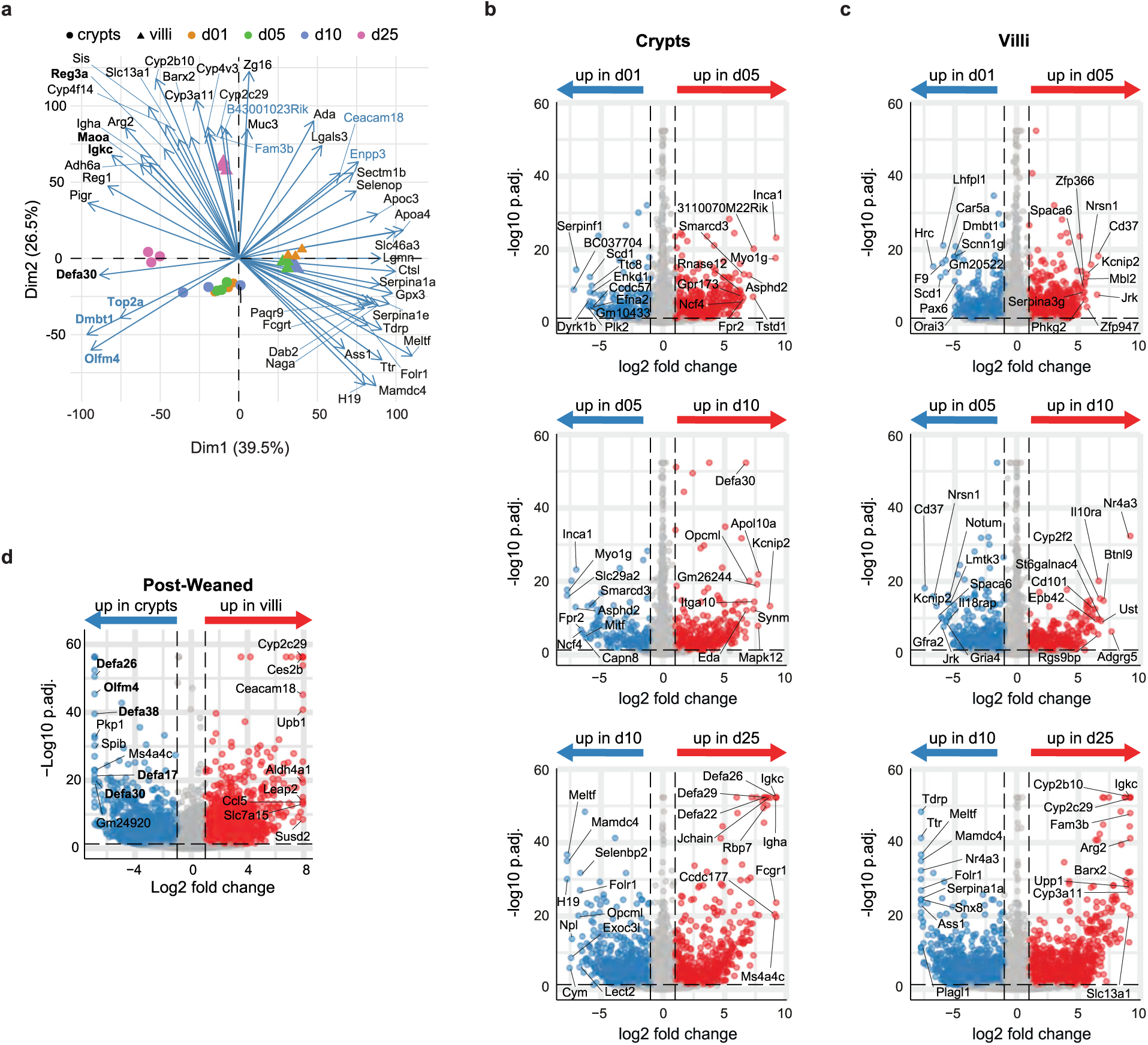
**(a)** Biplot displaying the top 50 driver genes for the 1^st^ and 2^nd^ dimension of the PCA plot shown in Fig. 3b. Black/blue names represent genes that were identified as DE/non-DE between pre- and post-weaning crypts or villi samples. 1-day-old (orange dots), 5-day-old (green dots), 10-day-old (blue dots) and 25-day-old (pink dots). **(b)** Volcano plots showing differential gene expression between 1- and 5-day-old (top), 5- and 10-day-old (middle) and 10- and 25-day-old (bottom) crypt samples. Blue and red dots indicate genes statistically (p_adj_<0.05) upregulated (|log2FC| > 1) in the younger and older animals, respectively. Grey dots indicate non-significantly and/or non-differentially regulated genes. **(c)** Volcano plots showing differential gene expression between 1- and 5-day-old (top), 5- and 10-day-old (middle) and 10- and 25-day-old (bottom) villi samples. Blue and red dots indicate genes statistically (p_adj_<0.05) upregulated (|log2FC| > 1) in the younger and older animals, respectively. Grey dots indicate non-significantly and/or non-differentially regulated genes. **(d)** Volcano plots showing differential gene expression between post-weaning (d25) crypt and villus samples. Blue and red dots indicate genes statistically (p_adj_<0.05) upregulated (|log2FC|>1) in the crypt and villi, respectively. Grey dots indicate non-significantly and/or non-differentially regulated genes.

**Supplementary Figure 4:**
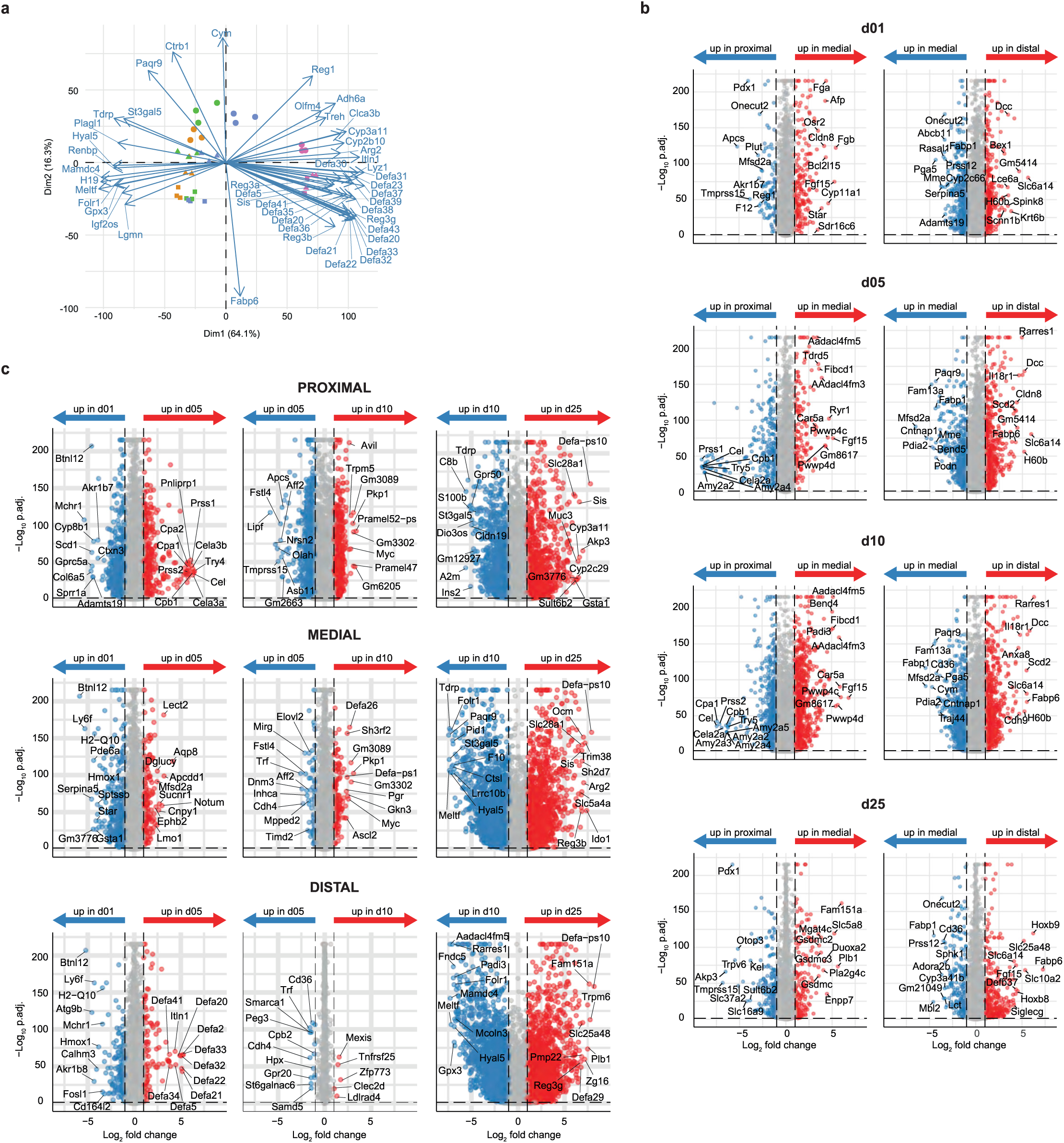
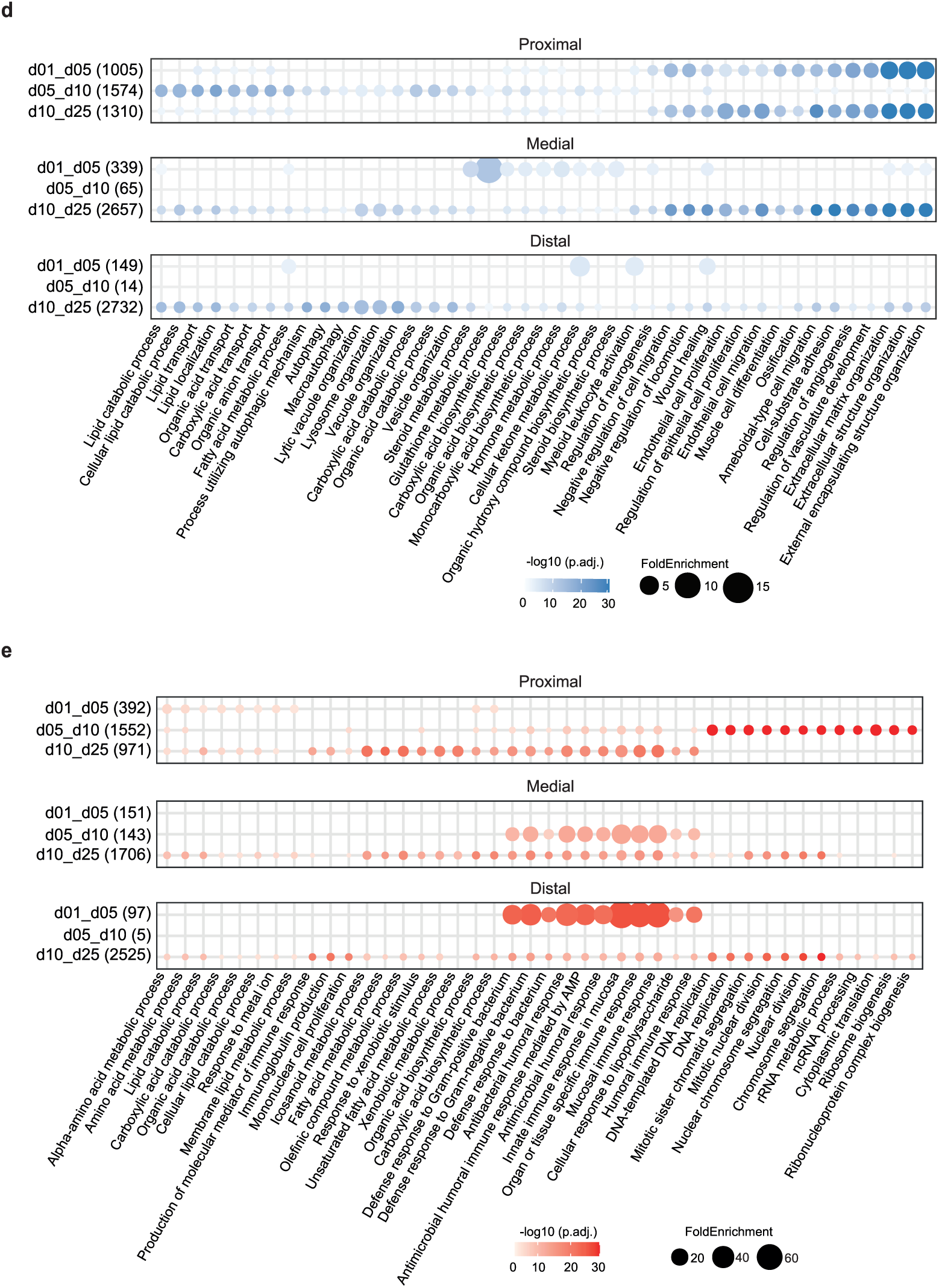
**(a)** Biplot displaying the top 50 driver genes for the 1^st^ and 2^nd^ dimension of the PCA plot shown in Fig. 4b. 1-day-old (orange dots), 5-day-old (green dots), 10-day-old (blue dots) and 25-day-old (pink dots). **(b)** Volcano plots showing differential gene expression between proximal and medial (left) and medial and distal (right) of 1-, 5-, 10 and 25-day-old animals (top to bottom). Blue and red dots indicate genes statistically (p_adj_<0.05) upregulated (|log2FC| > 1) in the most proximal and most distal parts, respectively. Grey dots indicate non-significantly and/or non-differentially regulated genes. **(c)** Volcano plots showing differential gene expression between 1- and 5-day-old (left), 5- and 10-day-old (middle) and 10- and 25-day-old (right) proximal, medial and distal samples (top to bottom). Blue and red dots indicate genes statistically (p_adj_<0.05) upregulated (|log2FC|>1) in the younger and older animals, respectively. Grey dots indicate non-significantly and/or non-differentially regulated genes. **(d)** Dot plot showing the top 10 enriched GO terms (by adjusted p-value) for the differentially expressed genes downregulated with age in the proximal, medial and distal parts (top to bottom). The number of DE genes associated with a GO term for each comparison is indicated in brackets. **(f)** Dot plot showing the top 10 enriched GO terms (by adjusted p-value) for the differentially expressed genes upregulated with age in the proximal, medial and distal parts (top to bottom). The number of DE genes associated with a GO term for each comparison is indicated in brackets.

**Supplementary Figure 5:**
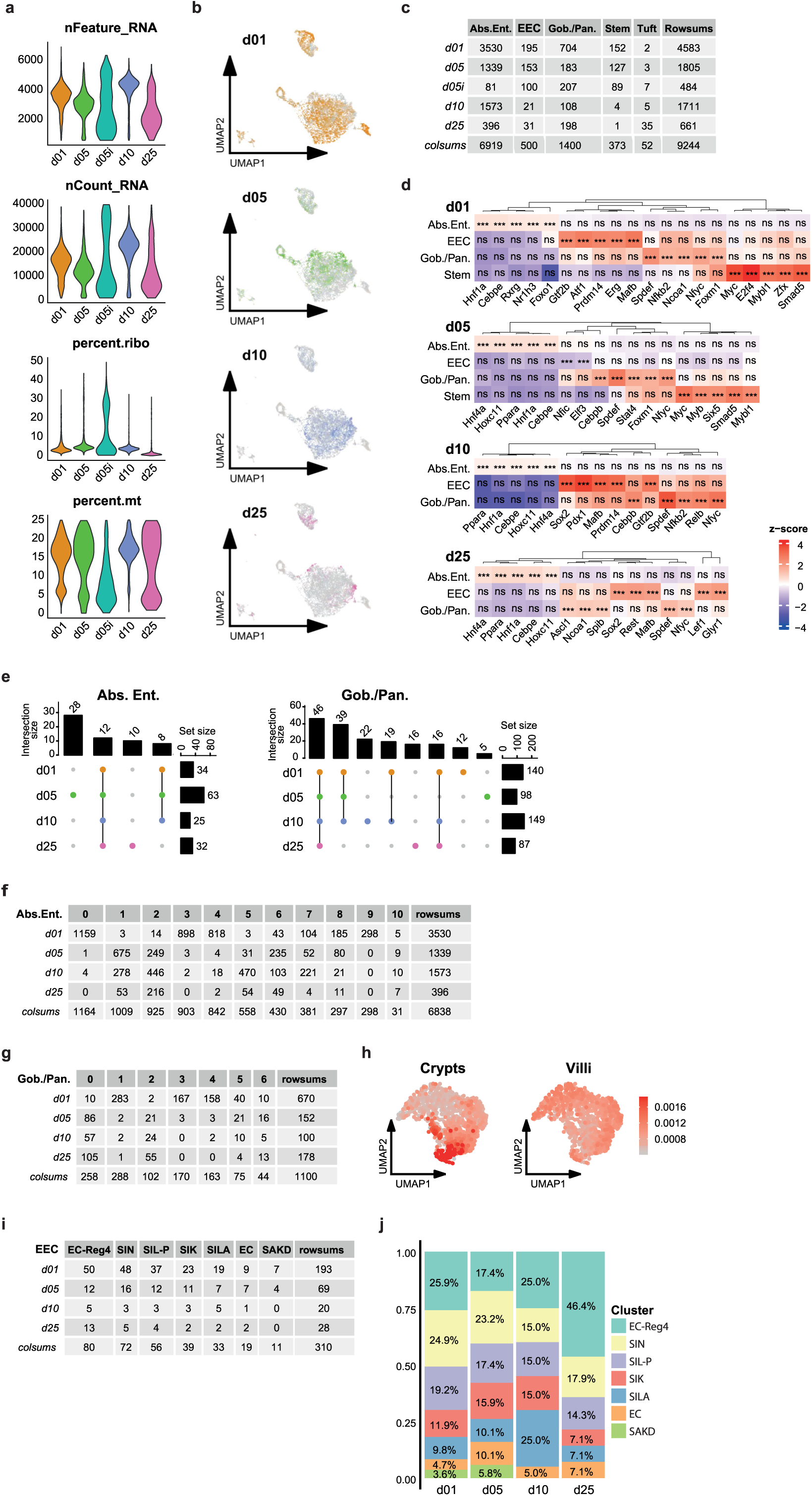
**(a)** Violin plots showing the feature counts, gene counts, and percentage of ribosomal and mitochondria genes after filtering of the scRNA-Seq dataset. Uninfected 1-day-old, orange; uninfected 5-day-old, green; 5-day-old infected at birth with *S.* Typhimurium, turquoise, uninfected 10-day-old, blue; uninfected 25-day-old, pink. **(b)** UMAP plot displaying the cells recovered at each timepoint. Same colour code as in Fig. S5a **(c)** Number of cells recovered per cell type and per time point after filtering. **(d)** Heatmap depicting the top 5 transcription factors (TF) per cell type predicted by DoRothEA in 1-, 5-, 10- and 25-day old animals. **(e)** Upset plots showing the number of DoRothEA-predicted transcription factors specific to one or more age group for the absorptive enterocyte (left) and Goblet/Paneth clusters (right). Interactions with less than 5 TFs are not shown. **(f)** Number of cells per absorptive enterocyte subcluster and per time point. **(g)** Number of cells per Goblet/Paneth subcluster and per time point. **(h)** UMAP plots showing the expression level of genes previously found to be associated with pre-weaned crypts (top left) and pre-weaned villi (bottom left), and the different subclusters recovered from the Goblet/Paneth cluster. The red colour intensity of the left plots reflects the overall gene expression level. **(i)** Number of cells per EEC subcluster and per time point. **(j)** Frequency of the 7 subtypes of EECs per time point. Subcluster colour code as in Fig. 5j.

**Supplementary Figure 6:**
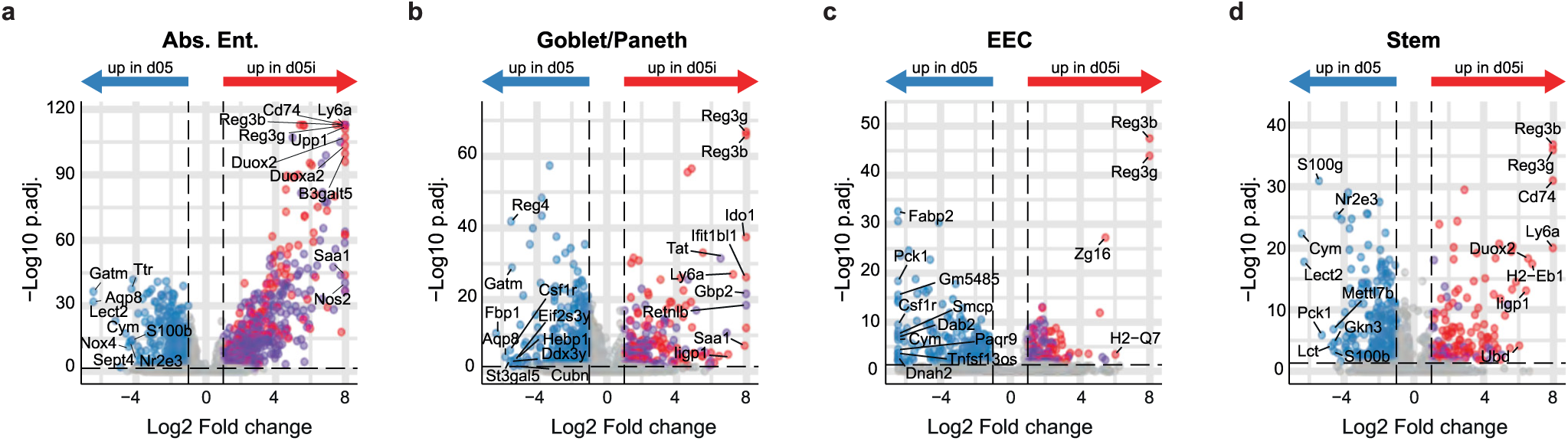
**(a-d)** Volcano plots showing differential gene expression between 5-day-old uninfected and *Salmonella*-infected (a) absorptive enterocytes, (b) Goblet/Paneth, (c) EECs and (d) stem cells. Blue and red/purple dots indicate genes statistically (p_adj_<0.05) upregulated (|log2FC| > 1) in the uninfected and infected samples, respectively. Purple dots indicate genes uniquely differentially regulated upon infection by one cell type, whereas red dots indicate gene differentially regulated by multiple cell types. Grey dots indicate non-significantly and/or non-differentially regulated genes.

